# The H3.3 K36M oncohistone disrupts the establishment of epigenetic memory through loss of DNA methylation

**DOI:** 10.1101/2023.10.13.562147

**Authors:** Joydeb Sinha, Jan F. Nickels, Abby R. Thurm, Connor H. Ludwig, Bella N. Archibald, Michaela M. Hinks, Jun Wan, Dong Fang, Lacramioara Bintu

## Abstract

Histone H3.3 is frequently mutated in cancers, with the lysine 36 to methionine mutation (K36M) being a hallmark of chondroblastomas. While it is known that H3.3K36M changes the cellular epigenetic landscape, it remains unclear how it affects the dynamics of gene expression. Here, we use a synthetic reporter to measure the effect of H3.3K36M on silencing and epigenetic memory after recruitment of KRAB: a member of the largest class of human repressors, commonly used in synthetic biology, and associated with H3K9me3. We find that H3.3K36M, which decreases H3K36 methylation, leads to a decrease in epigenetic memory and promoter methylation weeks after KRAB release. We propose a new model for establishment and maintenance of epigenetic memory, where H3K36 methylation is necessary to convert H3K9me3 domains into DNA methylation for stable epigenetic memory. Our quantitative model can inform oncogenic mechanisms and guide development of epigenetic editing tools.

## Introduction

Epigenetic memory, or the faithful maintenance of epigenetic information across DNA replication and cell division, is crucial for sustaining cell identity and tissue homeostasis. In multicellular organisms, maintenance of the epigenome relies on precise and coordinated regulation of gene expression, chiefly through control of DNA methylation and post translational modifications (PTMs) of histones.^1–3^ Loss of epigenetic information has been commonly linked to diseases such as accelerated aging and predisposition to types of cancers. ^4,5^ Indeed, somatic mutations of histone H3 and its variants, especially at regions in close proximity to PTM sites, are recurrent drivers in various human cancers. For example, studies identified lysine27-to-methionine (K27M) as the driver of diffuse midline glioma, while glycine 34 to arginine/tryptophan (G34 R/W) was found to be associated with glioblastoma and giant cell bone tumors, respectively.^6–8^ These somatic histone mutations, collectively referred to as oncohistones, often disrupt interactions between nucleosomes and their cognate chromatin readers, resulting in dysregulation of the epigenome and driving cells toward oncogenesis.

H3.3 K36M is an oncohistone found as a driver of various cancers and is prevalent in 95% of chondroblastoma cases. It involves a lysine 36 to methionine mutation of the *H3F3B* gene. The K36M oncohistone is a dominant negative mutant, meaning a single copy of the gene is sufficient for oncogenesis. H3.3 K36M has been shown to inhibit H3K36 methyltransferases SETD2 and NSD2 ^9,10^ This results in a global loss of both H3K36me2 and H3K36me3 and a redistribution of PRC2 catalyzed H3K27me3. Recent work has suggested that the H3.3 K36M oncohistone also acts in cis by inhibiting H3K36 methylation at genes where it incorporates, leading to derepression of transposable elements in drosophila. ^11,12^

H3K36 methylation is a ubiquitous histone modification conserved across eukaryotes and is prevalent in yeast, drosophila and humans. H3K36 can be mono, di or tri methylated and is associated with diverse functions including DNA repair, transcriptional elongation, and pre-mRNA splicing.^13,14^ Whereas H3K36me3 is catalyzed by a single methyltransferase, SETD2, H3K36me2 can be deposited by multiple enzymes, among them SETD2 and NSD1/2/3.^15,16^ Misregulation NSD1/2/3 or SETD2 expression have been commonly associated with cancers, suggesting epigenetic H3K36 methylation is important in oncogenesis. Although H3K36me2/3 is typically considered a hallmark of active transcription elongation, recent studies reported the presence of H3K36me3 in heterochromatin domains ^17^ and its contribution to inhibiting cryptic transcription. ^13,15^ Moreover, H3K36me2 and H3K36me3 is specifically recognized by the PWWP domains of de novo DNA methylases DNMT3A and DNMT3B ^20,71,72^, enzymes generally associated with gene silencing ^23,33,34^. These connections of H3K36me3 with heterochromatin and DNA methylation suggest a potential role in gene silencing or epigenetic memory in mammals. ^17–20^ However, mechanisms by which H3K36 methylation dynamically controls gene repression by interacting with heterochromatin is less well understood.

Constitutive heterochromatin is characterized by compact, transcriptionally repressed regions of the genome, canonically defined by the presence of H3K9me3. ^21–23^ Deposition of H3K9me3 is catalyzed by SETDB1 or SUV39H1/2, which convert H3 mono and dimethylation into trimethylation. This contributes to chromatin compaction, transcriptional repression, and a reduction of histone acetylation. ^24,25^ Furthermore, recruitment of these proteins leads to propagation of heterochromatin domains over tens to hundreds of kilobases. ^24,26–28^ This spreading occurs through reader writer-feedback: chromodomain-containing proteins such as HP1 bind to H3K9me3 and recruit more H3K9 methyltransferases, either directly or indirectly. When coupled with DNA methylation maintenance complexes consisting of DNMT1 and UHRF1, H3K9me3 can enable stable heritable gene silencing. ^23,29–32^

DNA methylation is a hallmark of epigenetic memory, as it is maintained robustly through cell division and is crucial in processes such as development and differentiation. ^1,23,34,35^ DNA methylation and H3K9me3 are both often found in regions of heterochromatin, suggesting an intimate connection between these epigenetic marks for gene silencing. ^36,37^ However, studies in lower eukaryotes have shown that deposition of DNA methylation usually depends on preexisting H3K9me3, suggesting H3K9me3 recruits proteins which write DNA methylation to reinforce gene silencing. ^38,39^ While detailed mechanisms of mammalian recruitment of DNMT3A/B still remain unclear, evidence suggests that this is conserved in higher eukaryotes as well: H3K9me3-associated proteins HP1 and SUV39H1 have been shown to interact with DNMT3A/B in mice and humans. ^40,41^ Given the prevalent occurrence of DNA hypomethylation in tumors and recurring dependence of SETDB1 expression in various cancer types, heterochromatin remains a therapeutic opportunity for clinical intervention in oncology. ^42–44^

To better understand how H3K36 methylation interacts with heterochromatin to regulate gene repression and epigenetic memory, as well as how this may be dysregulated in oncohistone pathophysiology, we develop a synthetic reporter system to nucleate heterochromatin by targeted recruitment of the repressive KRAB domain of ZNF10. ^26,45^ (KRAB is known to interact with KAP1, which in turn recruits SETDB1 and HP1, similar to endogenous heterochromatin formation. ^46,47^ We use this system to investigate the interplay between heterochromatin-mediated silencing and memory (during KRAB recruitment and release respectively) and H3K36 methylation in the context of H3.3 K36M expression, and find that oncohistone incorporation inhibits establishment of epigenetic memory in both HEK293T cells and a chondroblastoma cell culture model. We show these effects occur through changes in DNA methylation as a result of reduced H3K36 methyltransferase activity. We use these data to develop a 3-state stochastic model of heterochromatin spreading which accurately recapitulates and predicts epigenetic memory landscapes for normal cells as well as the H3.3 K36M disease state.

## Results

### H3.3 K36M disrupts KRAB mediated establishment and maintenance of epigenetic memory in HEK293T cells

To measure and monitor changes in gene expression upon H3.3 K36M expression, we engineered HEK293T cells with a synthetic transgene reporter^48^ integrated into the AAVS1 safe harbor locus using homology-directed repair. The reporter contains 9 copies of the tet operator site (TetO) upstream of a constitutive pEF promoter driving the expression of Citrine coupled to an IgG surface marker **(Fig. 1A)**. These cells also stably express the commonly used KRAB repressor domain from ZNF10 fused to the reverse tetracycline repressor (rTetR), allowing for targeted doxycycline (dox) dependent recruitment and subsequent long-term silencing of the reporter through deposition of H3K9me3 mediated heterochromatin.^28,45,49–52^

**Fig. 1.**
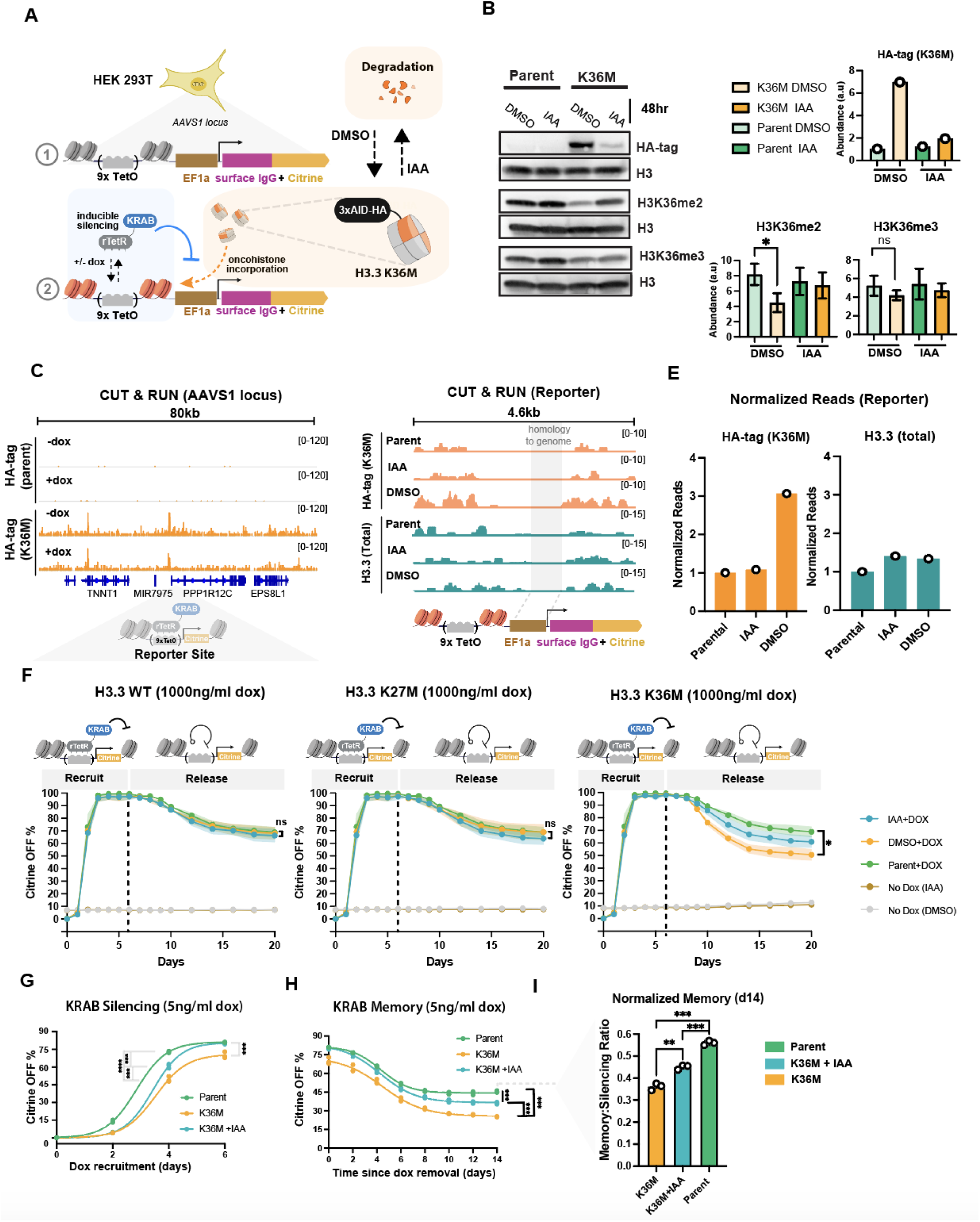
H3.3 K36M disrupts KRAB mediated establishment and maintenance of epigenetic memory in HEK23T cells. **(A)** Schematic of the synthetic constitutive reporter integrated in the AAVS1 locus of HEK293T, to which rTetR-KRAB can be recruited with doxycycline (dox) in the presence or absence of oncohistone incorporation to induce gene silencing. The reporter consists of 9 copies of the TetO binding site upstream of an EF1A promoter and drives the expression of an IgG surface marker (used for magnetic cell separation) and an mCitrine fluorescent protein. The oncohistone is tagged with the AID degron allowing for auxin (IAA) inducible degradation of H3.3 K36M. and an HA tag for detection. **(B)** Representative western blots with antibodies against HA-tag (for oncohistone expression), as well as histone modifications H3K36me2 and H3K36me3 in parent cells that express the reporter and KRAB but not the oncohistone, and in K36M cells that have all three components: reporter, KRAB and H3.3K36M.IAA was removed for 48 hours and replaced with carrier (DMSO) to allow accumulation of the oncohistone in the K36M. On the right, bar plots indicate abundance as measured by Western blot band intensities with each antibody, normalized to H3 input (data presented as mean +/- STD of n=3 biological replicates). **(C)** CUT&RUN genome tracks measuring HA-tag signal from H3.3 K36M incorporation within an 80kb domain around the reporter in parental and H3.3 K36M cell lines. Measurements were made in the presence or absence of dox (1000 ng/ml) recruitment of KRAB for 6 days and total reads were normalized using E.coli DNA spike-in. **(D)** CUT&RUN genome tracks measuring incorporation levels of H3.3 K36M and total histone H3.3 (measuring both endogenous H3.3 and H3.3 K36M) at the AAVS1 reporter in the presence or absence of IAA induced degradation. Values are normalized by total number of reads and shown as counts per million. The EF1a promoter region is shaded gray and not analyzed since it is identical in sequence with the endogenous EF1a promoters. **(E)** Quantification of normalized reads at the reporter for CUT&RUN genome tracks shown in D. **(F)** Flow cytometry time courses in HEK293T measuring the mean fraction of Citrine OFF cells upon 6 days of 1000 ng/ml dox recruitment of KRAB and 14 days of memory. Shading represents SEM (n= 3 biological replicates). Dashed line indicates the time of dox removal. Statistical analysis was performed using Welch’s T-test. **(G)** Flow cytometry time course of low-recruitment KRAB silencing in the absence of IAA (no degradation) in cells expressing H3.3 K36M (orange) and parental cells without the oncohistone (green) at low levels of dox (5ng/ml dox). IAA (50μM) is added (blue curve) to partially degrade H3.3 K36M. n= 3 biological replicates. Statistical analysis was performed using a one-way ANOVA. **(H)** KRAB memory time course: cells were first silenced at low dox (5ng/ml, as shown in G), then dox was completely removed and the percentage of cells with the citrine reporter off is plotted for 14 days. Same colors and analyses as in panel G.**(I)** Bar plots quantifying memory data in H where the % of Citrine OFF is normalized at day 14 to the level of each cell line’s level of silencing at day 0 of dox removal (day 6 of silencing). n= 3 biological replicates. Statistical analysis was performed using a one-way ANOVA

To enable conditional expression of the oncohistone in the reporter cells, we stably expressed H3.3 K36M fused to an auxin inducible degron (AID, **Fig. 1A right**). This system allows auxin (IAA) dependent degradation^53^ of the oncohistone or oncohistone accumulation in the absence of IAA (DMSO carrier only), as detected by Western blotting against the HA tag also included in the oncohistone **(Fig. 1B)**. Addition of IAA leads to partial degradation of the oncohistone, allowing us to test phenotypes for intermediate oncohistone concentrations **(Fig. 1B right)**. This system can recapitulate epigenetic changes observed in other models of chondroblastoma: in the K36M line treated with DMSO (allowing oncohistone accumulation), expression of H3.3 K36M **(Fig. 1B left)** led to a loss of H3K36me2, and to a lesser degree, a loss of H3K36me3 **(Fig. 1B right)** compared to the parental line that does not express the oncohistone, consistent with previous studies.^10,11,54^

To verify incorporation of oncohistone into chromatin across the genome and at our reporter, we performed CUT&RUN ^55^ using antibodies against the HA-tag that is part of the oncohistone-degron fusion after 48 hours of oncohistone expression (IAA removal, DMSO only). We observe a high level of H3K36M oncohistone incorporation (HA-tag) both across the genome, localized mostly to promoters, exons and distal intergenic regions **(Fig. S1A&B)**, and at the AAVS1 reporter integration locus, both with and without KRAB recruitment (+/-dox) **(Fig. 1C)**. Moreover, H3K36M is incorporated in our reporter region, as shown by the higher number of reads in the presence of oncohistone compared to the parental line that does not express the oncohistone **(Fig. 1D&E, left)**. In contrast, total levels of reads from H3.3 CUT&RUN (which detects both WT H3.3 and the mutant H3.3) remain relatively unchanged (**Fig. 1E, right**).

Having established a reporter system that allows for heterochromatin generation at the AAVS1 locus simultaneously with local oncohistone incorporation, we asked whether H3.3 K36M has an effect on KRAB-mediated silencing or the ability of maintain stable silencing over multiple cell divisions after removal of dox (epigenetic memory). To quantify effects on silencing dynamics, we first removed IAA for 48hrs to allow oncohistone pre-incorporation, then added dox to recruit KRAB for 6 days and measured the percentage of silenced reporter cells using flow cytometry. Upon addition of saturating dox (1000 ng/ml), by the end of 6 days KRAB completely silences the reporter in nearly 100% of the cells across all cell lines expressing H3.3 variants: wild type (WT), K36M and K27M. **(Fig. 1F).** After KRAB release (dox removal), the H3.3 K36M cells showed reduced memory, reactivating the reporter to a higher degree than the parental line (**Fig. 1F, right,** DMSO vs parental). We did not observe this decrease in memory upon expression of WT H3.3 or another oncohistone, H3.3K27M. Furthermore, by adding IAA to induce partial degradation of the H3.3 K36M oncohistone **(Fig. 1B)**, we were able to partially rescue this reduced memory phenotype (**Fig. 1F, right, IAA)**. We verified that these effects of silencing and memory were not due to differential growth of the cell lines or aberrant expression of the basal reporter levels **(Fig. S1C,D & E).**

Additionally, to further interrogate whether observed incorporation of the oncohistone was necessary for observed effects on memory, we expressed a truncated version of the H3.3 K36M lacking the globular domain, preventing it from being loaded onto nucleosomes. This approach is similar to that used in a previous study. ^56^ Despite this construct containing the first 44 amino acids of H3.3 including the K36M tail, we observed no changes in epigenetic memory **(Fig. S1G).** This suggests full length H3.3 K36M causes decreased memory through its incorporation into chromatin.

Given H3K36 methylation is commonly associated with active genes ^11,13^, and that K36M increases the percentage of reactivated cells after KRAB release **(Fig. 1F)**, we also tested the effect of H3.3 K36M on gene activation from a minimal promoter in the presence of a commonly used transcriptional activator, VP64. ^57^ We observed no changes in gene activation upon low or high levels of dox-mediated VP64 recruitment **(Fig. S1F)**.

Given a significant loss of epigenetic memory in the H3.3 K36M cells occurred without a change in the level of silenced cells at day 6 of KRAB recruitment, we revisited the effects of silencing using lower concentrations of dox in case effects on silencing were being masked at saturating doses. When we repeated our KRAB recruitment assays at 5 ng/ml dox, we saw that the H3.3 K36M cell line exhibited slower silencing than the parental line **(Fig. 1G)**. We again noticed a similar effect on memory, where the H3.3 K36M cell line exhibited a lower level of memory after dox removal compared to parental cells **(Fig. 1H)**. We see this decrease in memory even when normalizing the final memory to account for differences in silencing between cell lines at the end of recruitment **(Fig. 1I).** Additionally, partial degradation of the oncohistone (by IAA addition) partially rescued the silencing and memory phenotypes (**Fig. 1G-I**), suggesting slower silencing and reduced memory at non-saturating KRAB recruitment is a direct result of the H3.3 K36M oncohistone expression as opposed to intrinsic differences between the cell lines.

### KRAB mediated epigenetic memory and silencing speed is reduced in a H3K36M chondroblastoma model

We next wanted to determine whether the effects we observe with H3.3 K36M on KRAB silencing and epigenetic memory in HEK293T and oncohistone overexpression can be recapitulated in a physiological disease model where an endogenous H3.3 gene contains the K36M mutation. To address this, we utilized TC28a2 immortalized chondrocytes due to their common use as an established model for studies of human bone and cartilage pathophysiology.^58,59^ In cell culture models, editing of an endogenous H3.3 allele to introduce the mutation in TC28a2 cells has been shown to be sufficient to recapitulate effects of the chondroblastoma, including differentiation defects and changes in gene expression level from RNAseq and epigenetic changes.^10^ We thus utilized cells from the same study where one copy of the endogenous histone H3.3 (*H3F3B* gene) was edited in two separate clones using CRISPR to introduce an endogenous lysine to methionine point mutation at lysine 36 in *H3F3B*. We then introduced our Citrine reporter into the AAVS1 locus into these cells using TALEN mediated DNA integration, as well as the rTetR-KRAB expression vector using lentivirus to enable dox mediated induction of gene silencing through targeted recruitment **(Fig. 2A)**. We verified that the knock-in mutations at the *H3F3B* gene were maintained in our reporter clones using Sanger sequencing **(Fig. S2A).** We also measured the levels of H3K36 methylation in each line using western blot and observed that both edited clones exhibited lower levels of H3K36me2 and H3K36me3 compared to the WT line **(Fig. 2B)**, consistent with the literature.^10^

**Fig. 2.**
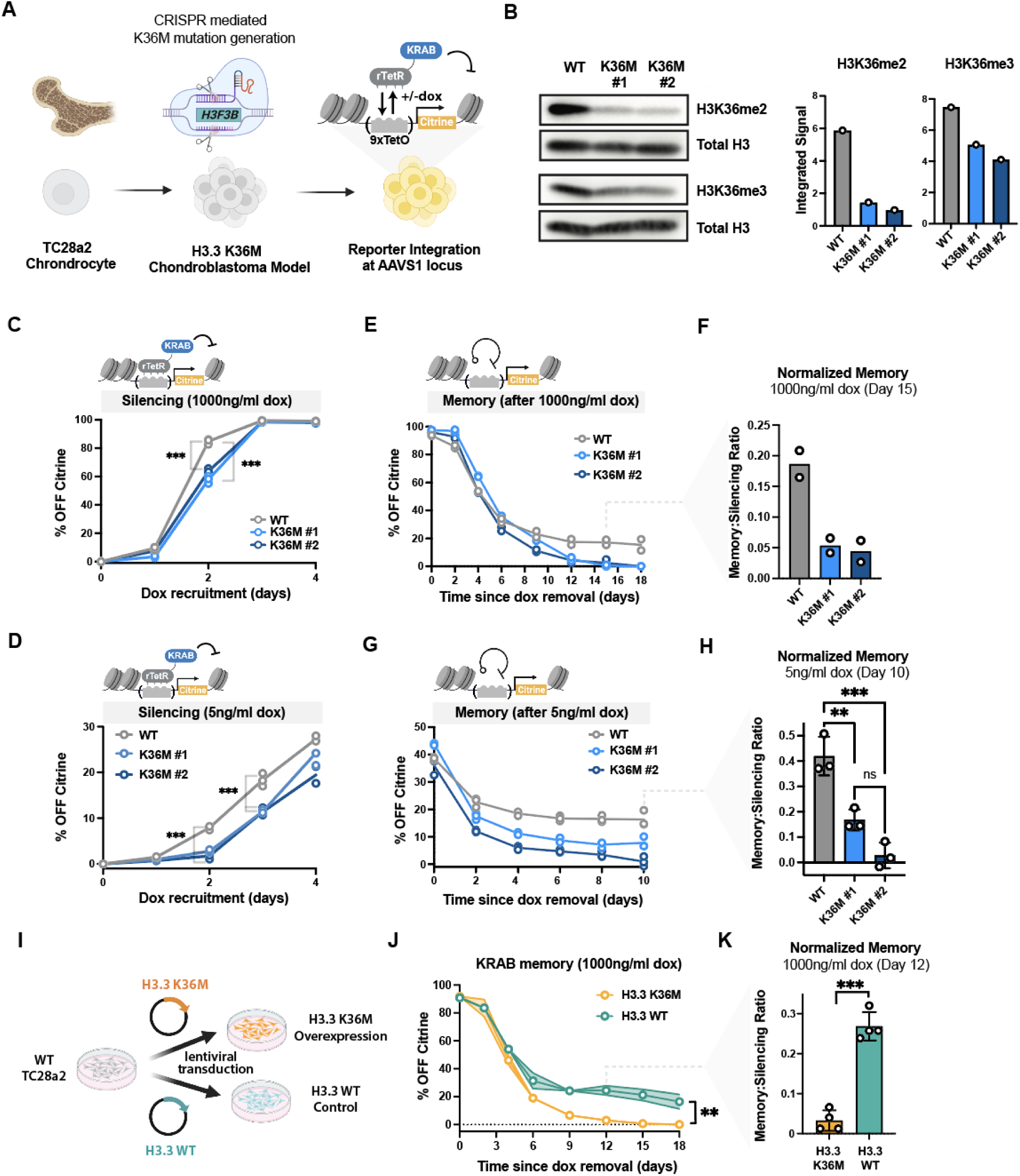
KRAB mediated epigenetic memory and silencing speed is reduced in a H3K36M chondroblastoma model. **(A)** Schematic describing the workflow to engineer TC28a2 chondrocytes to generate chondroblastoma reporter cell lines: one copy of the endogenous histone H3.3 (*H3F3B* gene) was edited in two separate clones using CRISPR to generate the K36M mutation^10^. We integrated the Citrine reporter into the AAVS1 locus using TALENs, and delivered TetR-KRAB using lentivirus. **(B)** Western blots with quantification of band intensities showing levels of H3K36me2 and H3K36me3 (normalized by Total H3) in wildtype TC28a2 cells and two CRISPR edited K36M clones. **(C)** Flow cytometry time course during 4 days of KRAB recruitment (with 1000ng/ml dox) in TC28a2 cells, with the fraction of Citrine silenced cells quantified over time for WT (gray) and H3.3 K36M TC28a2 reporter clones(n= 3 biological replicates). Statistical analysis was performed using a one-way ANOVA **(D)** Flow cytometry time course during 4 days of KRAB recruitment with low dox (with 5ng/ml dox) in TC28a2 cells. The fraction of Citrine silenced cells quantified over time for WT and H3.3 K36M TC28a2 reporter clones (n= 3 biological replicates). Statistical analysis was performed using a one-way ANOVA. **(E)** Flow cytometry time course measuring epigenetic memory in TC28a2 cells, shown as percentage of cells with citrine silenced after 5 days of KRAB recruitment (at 1000ng/ml dox) and release (dox removal) for 18 days. The fraction of Citrine silenced cells is measured after dox removal at day 0 and normalized to the no dox control at each time point (n=2 biological replicates, see methods). **(F)** Bar plots quantifying normalized epigenetic memory from the time-courses in E by dividing memory (the %Citrine OFF cells at day 15) to the initial silencing (%OFF at day 0) from n=2 biological replicates. **(G)** Flow cytometry time course measuring epigenetic memory after 6 days of KRAB recruitment (5ng/ml dox) in TC28a2 cells. The fraction of Citrine silenced cells is measured after dox removal at day 0 and normalized to the no dox control at each time point (see methods). n=3 biological replicates. Statistical analysis was performed using a one-way ANOVA. **(H)** Bar plots quantifying normalized epigenetic memory as a memory-to-silencing ratio from the time-courses in G by normalizing the %Citrine OFF cells at day 15 to the initial %OFF at day 0 (n=3 biological replicates). Statistical analysis was performed using a one-way ANOVA. **(I)** Schematic describing workflow to overexpress H3.3 (WT) or H3.3 (K36M) in wildtype TC28a2 cells. **(J)** Epigenetic memory time course after 5 days of KRAB recruitment (1000ng/ml dox) and 18 days of dox release in WT TC28a2 cells transduced with lentivirus consisting of H3.3 or H3.3 K36M overexpression. Shading represents standard deviation at each timepoint (n=4 biological replicates). Statistical analysis was performed using Welch’s T-test. **(K)** Bar plots quantifying normalized epigenetic memory calculated from the time-courses in J by normalizing memory (the %Citrine OFF cells at day 12) to silencing (the initial %OFF at day 0) from n=4 biological replicates. Statistical analysis was performed using Welch’s T-test.

To measure the effect of endogenous H3.3 K36M on silencing dynamics in the TC28a2 reporter lines, we recruited KRAB to the Citrine reporter for 5 days and measured the fraction of Citrine OFF using flow cytometry as before. At saturating dox, both mutant clones or the WT line silenced to nearly 100% OFF within 4 days **(Fig. 2C)**, similarly to HEK293T. However, both K36M clones had a small but significant reduction in silencing speed, as evident by a reduction of %OFF Citrine at day 2 compared to WT cells **(Fig. 2C)**. Moreover, as with HEK293T, lower strength recruitment of KRAB (lower dox) in TC28a2 cells led to slower silencing dynamics in the K36M oncohistone background compared to the wild-type. **(Fig. 2D).** We verified there was no appreciable difference between growth rate between the cell lines using a dye-spike in growth assay, eliminating dilution of the Citrine as a confounding reason for differences in silencing (**Fig. S2B, S2C**). Furthermore, the mean basal reporter expression in the K36KM and WT cells are within similar levels in the citrine-on population when compared to the magnitude of change induced by KRAB silencing (**Fig. S2D)**.

To measure effects on epigenetic memory, dox was removed after 5 days of KRAB recruitment, and reporter reactivation was assessed every two days with flow cytometry. We observed that both H3.3 K36M edited clones began to reactivate the reporter much more quickly than the WT line and stabilized at an irreversible memory fraction which was around 15% lower than the parental line. **(Fig. 2E&F)**. Additionally, we tracked the reactivation of these cells after low dox recruitment and, similarly to release after high dox, memory is higher in the WT cell line compared to both K36M clones **(Fig. 2G&H)**. This reactivation phenotype is similar to that observed in HEK293T cells expressing the exogenous H3.3 K36M, suggesting establishment of epigenetic memory is inhibited by the expression of the oncohistone.

Since H3.3 K36M overexpression is sufficient to transform phenotypically normal TC28a2 cells to become chondroblastoma-like or contribute to pathologies in human, mice and drosophila ^10–12,60^, we asked whether overexpression of the oncohistone in wildtype cells (as opposed to editing the endogenous H3.3 gene) is also associated with a loss in epigenetic memory. Given that wildtype TC28a2 cells exhibited robust epigenetic memory in earlier experiments, loss of memory would confirm this phenotype is a consequence of H3.3 K36M presence as opposed to a result of cell-cell heterogeneity associated with single-clones. We thus delivered the same oncohistone lentiviral vectors used in the HEK293T experiments (**Fig.1)** to overexpress H3.3 K36M or H3.3 WT in the WT TC28a2 reporter lines that exhibited epigenetic memory after KRAB recruitment (**Fig. 2I**). While overexpression of the WT H3.3 had no effects on epigenetic memory, with cells maintaining close to 20% of irreversible Citrine OFF cells after 5 days of KRAB recruitment, H3.3 K36M overexpression abolished nearly all memory **(Fig. 2J&K).** H3.3K36M overexpression was thus sufficient to reduce epigenetic memory in the TC28a2 cells to levels comparable to the K36M edited clones. Furthermore, these effects are specific to the H3.3 K36M oncohistone and are unlikely to be a result of intrinsic cell-cell heterogeneity, as expression of the wild type H3.3 did not have any appreciable effect on memory.

### Loss of epigenetic memory by H3.3 K36M is not due to reduced H3K9me3 deposition

Given differences in epigenetic memory observed between the WT and K36M cells, we wanted to confirm that individual cells in the population can maintain epigenetic memory across cell divisions, i.e. remain stably silenced when sorted in the memory phase. Given the fraction of irreversibly silenced cells by KRAB is dependent on the duration of recruitment ^50^, we reasoned that increasing recruitment time in the K36M cells would allow for higher levels of memory that would more closely resemble the parental line and enable more similar sort conditions. To this end, we silenced the reporter in HEK293T parental and K36M cells by treating them with dox for 2 days and 10 days respectively, to ensure the two cell lines reached roughly similar levels of cells silenced at the memory timepoint (14 days post dox removal, **Fig. 3A, pre-sort**). We then sorted the reporter OFF population (**Fig. 3A, Day 2)** using magnetic beads against the IgG cell surface marker included in our reporter ^48^ **(see Fig. 1A)**, and measured the fluorescence of these sorted OFF cells after multiple cell divisions **(Fig. 3A, Day 20)**. The final fluorescence distribution of these cells sorted once memory has stabilized remained relatively unchanged over the course of the following 18 days, suggesting that KRAB induced epigenetic memory is stable and irreversible at the single-cell level over our experimental timescales in both the WT and K36M cells.

**Fig. 3.**
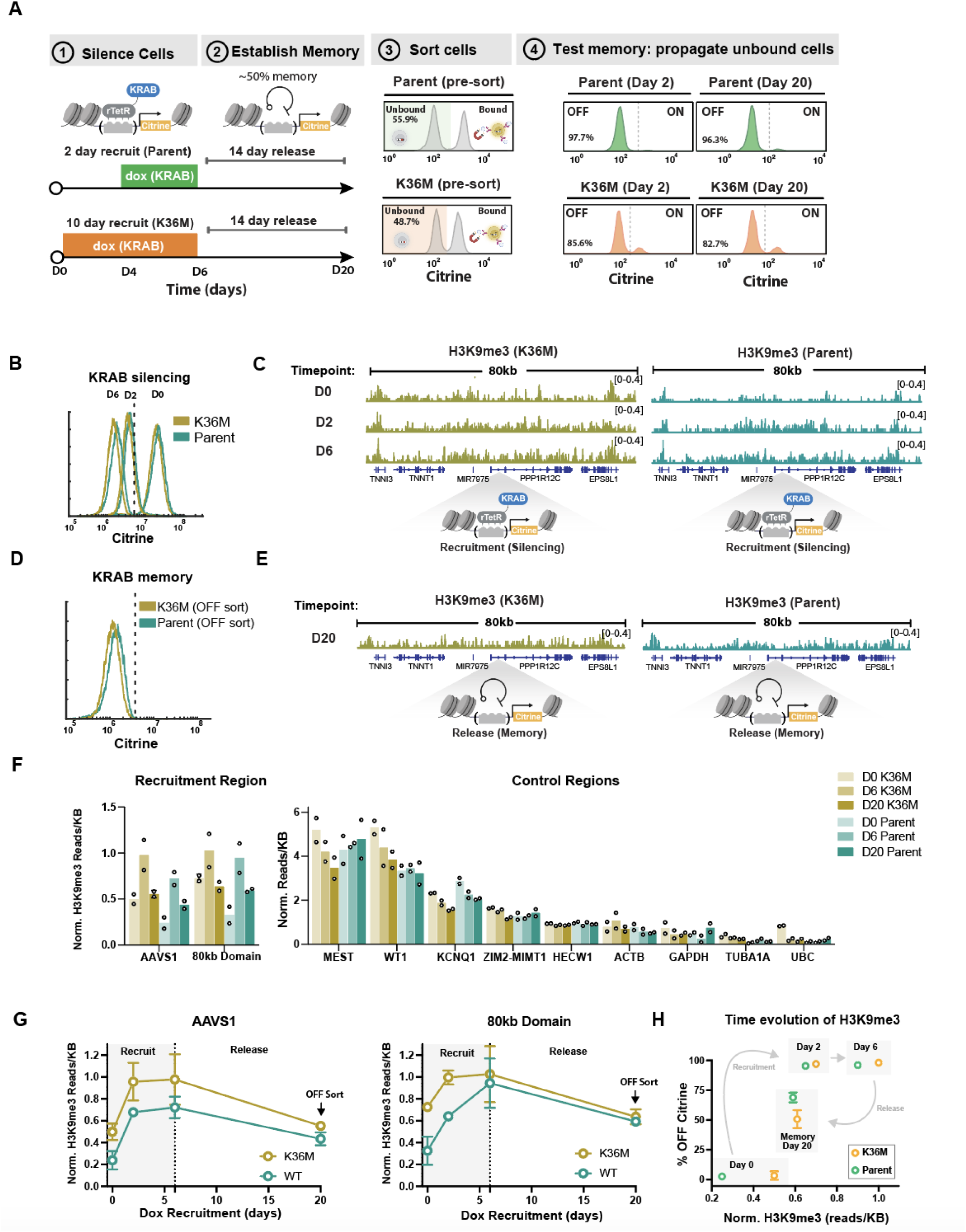
Loss of epigenetic memory by H3.3 K36M is not due to reduced H3K9me3 deposition. **(A)** Schematic describing magnetic cell separation after establishment of epigenetic memory. KRAB was recruited with dox (1000 ng/ml) for 2 days in parental cells and 10 days in K36M cells to generate roughly a 50% OFF memory population after 14 days of release. This OFF population was then sorted for each cell line, and silenced cells were propagated in cell culture to assess maintenance of memory using flow cytometry 2 days and 20 days after sorting. **(B)** Flow cytometry distributions of Citrine reporter expression during KRAB recruitment in parental and K36M cells for 0 days (no dox, D0), 2 days (D2) and 6 days (D6) with 1000 ng/ml dox. **(C)** CUT&RUN genome tracks for H3K9me3 during KRAB silencing for the same populations of cells shown in B. Read numbers are normalized by total number of reads and shown as counts per million. **(D)** Flow cytometry distributions of Citrine expression after enriching the stably silenced memory population by FACS 14 days after KRAB release. **(E)** CUT & RUN genome tracks for H3K9me3, normalized as in C, during KRAB memory corresponding to the sorted population of cells shown in D. **(F)** Left: H3K9me3 reads densities at the reporter (AAVS1) and the surrounding 80kb domain in the vicinity of the *PPP1R12C* gene quantified from genome tracks at the timepoints shown in C&E (n=2 biological replicates). Right: H3K9me3 reads densities from a panel of control genes (n=2 biological replicates). **(G)** H3K9me3 levels (counts per million per kb) at the AAVS1 locus (*PPP1R12C gene*) and the entire 80kb domain around the reporter over time in WT and K36M cells upon dox recruitment and release. The release time point is from the sorted memory population in D (n=2 biological replicates). **(H)** A sequential scatter plot of the dynamics of H3K9me3 versus the fraction of Citrine silent cells from flow cytometry data in Fig. 1F, showing the time evolution of the system in parental (green) and K36M cells (orange) for 0 days, 2 days and 6 days of silencing respectively. The Day 20 memory time point shows H3K9me3 levels for the sorted OFF cells in Fig. D. Percentages of cells OFF (Y-axis) are the mean of 3 biological replicates +/- SEM. H3K9me3 levels (X-axis) are the mean of 2 biological replicates (CUT & RUN from 3C&E).

We next sought to determine whether the reduced epigenetic memory observed with H3.3 K36M was associated with changes in KRAB mediated heterochromatin formation. One common hallmark of heterochromatin is stable and heritable gene silencing due to H3K9me3 domains that are maintained by positive feedback mechanisms.^61–63^ Therefore, we performed CUT&RUN ^64^ to measure levels of H3K9me3 at and around the reporter at different times after rTetR-KRAB recruitment. The flow cytometry distributions of Citrine expression were nearly indistinguishable between the WT and K36M cells, with nearly 100% of cells silenced in both cell lines **(Fig. 3B).** In both WT and oncohistone lines, recruitment of KRAB led to H3K9me3 deposition across a large region (∼80kb) across the AAVS1 locus **(Fig. 3C, Fig. S3A)**, similarly to studies performed in different cell lines. ^28^ This dox-mediated increase of H3K9me3 was also associated with a decrease of H3K4me3 at the AAVS1 locus **(Fig. S3 B&C)**, as expected from the reduction in transcription in both cell lines upon KRAB recruitment as well as retention of much of the H3.3K6M at the reporter despite H3K9me3 deposition **(Fig. S3 D&E)**.

Another common modification associated with heterochromatin and epigenetic memory is H3K27me3, deposited by the PRC2 complex. ^65,66^ Despite global increase in some control genes occurring in the K36M cell line compared **(Fig. S3H)**, likely due to a redistribution of PRC2 upon loss of H3K36 methylation ^67–70^, we saw no dox-mediated changes of H3K27me3 at the reporter region **(Fig. S3 F&G),** suggesting H3K27me3 is not involved in the H3K36M-induced changes in memory at our reporter.

We next sorted irreversibly silenced cells 20 days after dox removal (**Fig. 3D, Fig. S3I**), and measured H3K9me3 levels in both WT and K36M cells **(Fig. 3E).** In both cell lines, H3K9me3 modifications went down to levels similar to no dox (D0, no KRAB recruitment) (**Fig. 3F**), suggesting H3K9me3 is not necessary for maintaining memory at our reporter. Changes in H3K9me3 were localized to the site of recruitment; we did not observe any global systematic increase of H3K9me3, as evidenced at a panel of selected control genes displaying a range of H3K9me3 levels, as well as a subset of active housekeeping genes. **(Fig. 3F),** suggesting the observed changes at the reporter are due to KRAB-mediated silencing.

Quantification of the domain size over time during dox mediated recruitment and release reveals an increase of H3K9me3 at both the AAVS1 reporter and across the 80kb domain around the reporter insertion locus, which increases with KRAB recruitment time; however, the final H3K9me3 levels are similar in the WT and H3.3 K36M cells after 6 days of dox **(Fig. 3G).** Furthermore, 20 days after removal of dox, absolute H3K9me3 levels in the sorted irreversibly silenced cells fall back down to levels similar to one another in both the WT and H3.3 K36M cells, suggesting differences in epigenetic memory are not a result of different levels of H3K9me3. Consolidating our CUT & RUN data **(Fig. 3C&E)** and previously shown flow cytometry time-course data **(Fig. 1F)**, we tracked the time-evolution of H3K9me3 and Citrine reporter fluorescence during KRAB recruitment and release to visualize the dynamics of H3K9me3 and gene silencing (**Fig. 3H**): Cells start with high levels of Citrine and low levels of H3K9me3 at day 0 before recruitment of KRAB. As dox is added, H3K9me3 increases in both WT and H3.3 K36M cells along with a decrease of Citrine, signifying gene silencing. Within 2 days of KRAB recruitment, the Citrine is fully silenced, however, H3K9me3 continues to increase up to day 6 when measuring the 80kb domain around the reporter. This suggests spreading of the modification over these timescales in both the WT and the K36M cells. Upon removal of dox, a higher fraction of cells remain OFF in the WT population compared to H3K36M, though the sorted OFF cells have the same levels of H3K9me3 in the two cell lines **(Fig. 3H Memory)**, suggesting that a mechanism separate from H3K9me3 is responsible for the difference in epigenetic memory we observe with the oncohistone.

### H3.3 K36M disrupts the deposition of DNA methylation upon KRAB recruitment

Given the observed loss of epigenetic memory in the *H3F3B* edited TC28a2 cells and HEK293T cells exogenously expressing H3.3 K36M, we speculated that levels of H3K36 methylation may play a role in the stable maintenance of KRAB mediated gene silencing. Other groups have shown that H3.3 K36M can inhibit the methyltransferase activity of NSD2 and SETD2, which can result in a decrease of H3K36me2 and H3K36me3 across the genome.^10,54,60^ We thus asked if siRNA knockdown of methyltransferases involved in H3K36 methylation would be sufficient to observe changes in epigenetic memory and phenocopy the behavior of the oncohistone upon KRAB recruitment in wildtype HEK293T cells in the absence of any H3.3 K36M **(Fig. 4A)**. Knockdown of methyltransferase NSD1 using siRNA was sufficient to reduce both global levels of H3K36me2 and H3K36me3, while SETD2 knockdown only reduced levels of H3K36me3 **(Fig. 4B)**, as expected, since it catalyzes the H3K36me2 to me3 transition **(Fig. 4A)**. Similarly, NSD2 and NSD3 knockdown reduced H3K36me2, though to a lesser extent **(Fig. S4A)**. We verified that all the siRNAs used in isolation without recruitment of KRAB did not affect the basal reporter expression **(Fig. S4B).** Since NSD1 and SETD2 siRNAs had the strongest effects on H3K36 methylation, we tested them for effects on epigenetic memory.

**Fig. 4.**
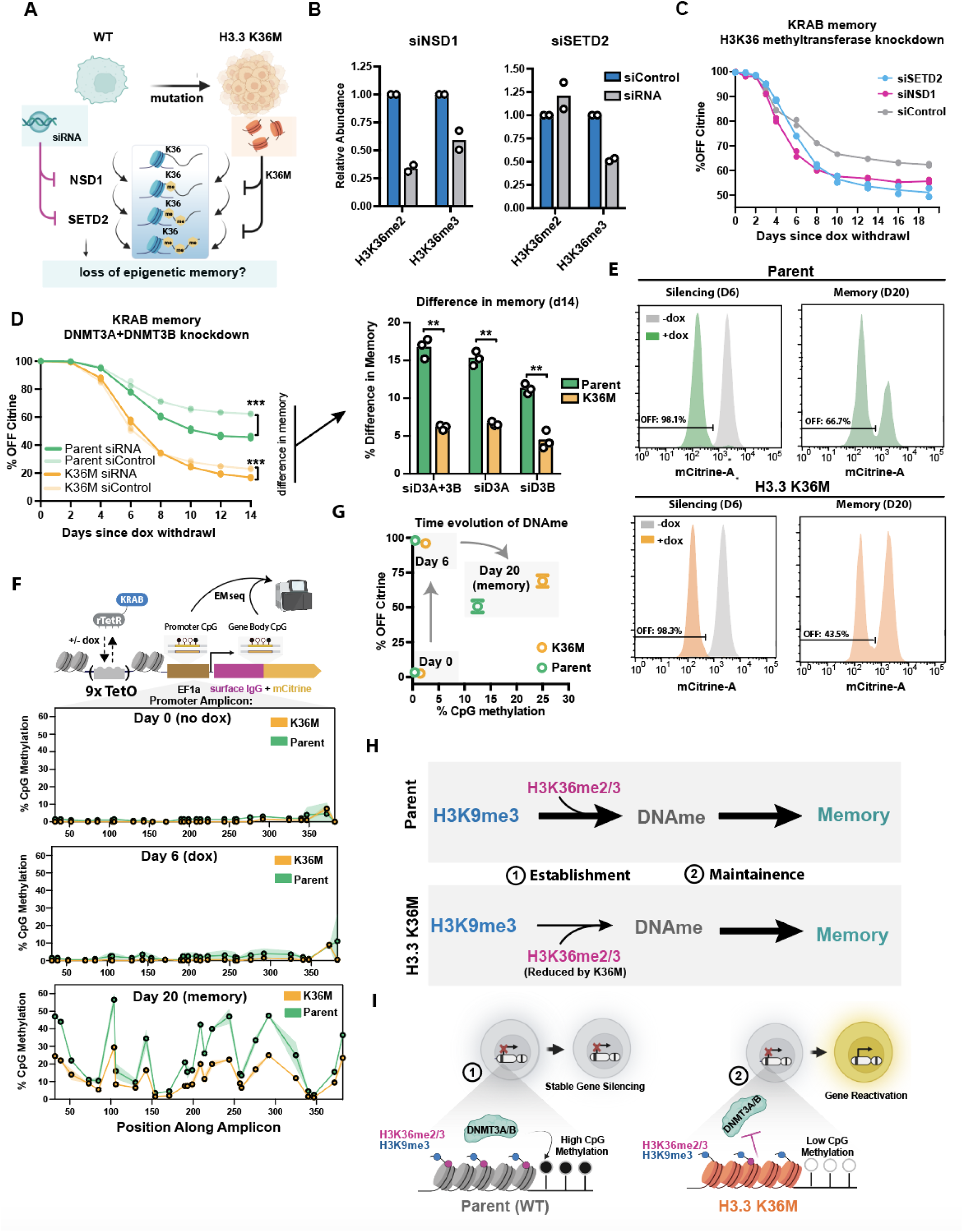
H3.3 K36M disrupts the deposition of DNA methylation after KRAB recruitment and release. **(A)** Schematic describing siRNA perturbations targeting H3K36 methyltransferases in wildtype cells to phenocopy effects of the H3.3 K36M oncohistone. **(B)** Quantification of global H3K36me2 and H3K36me3 levels by Western blot 48hr post-transfection with siRNA targeting NSD1 or SETD2 respectively (n= 2 biological replicates), normalized first to H3 levels for each condition, and then relative to siRNA control. **(C)** Flow cytometry time courses measuring epigenetic memory in HEK293T cells treated with siRNA targeting either NSD1 or SETD2 during the memory phase after KRAB recruitment (n = 2 biological replicates), with siRNA delivery at day 0. Cells were silenced by KRAB recruitment with 1000ng/ml dox for 2 days prior to beginning the time course shown. **(D)** Left: Flow cytometry time courses measuring epigenetic memory in HEK293T cells treated with siRNA targeting de novo methyltransferases DNMT3A and DNMT3B in WT (green) or H3.3 K36M (orange) cells (with siRNAs delivered at day 0 of memory, upon dox removal). Darker color lines represent targeting siRNA and lighter color lines represent non-targeting control siRNA in each cell line (n=3 biological replicates). The black line at day 14 indicates the difference in memory at the endpoint of the experiment. Cells were silenced by KRAB recruitment with 1000ng/ml dox for 2 days prior to beginning the dox removal time course shown. Right: Bar plots quantifying the difference in memory between the DNMT3A/B target siRNA and control siRNA in WT and H3.3 K36M cells at day 14 after dox removal (n=3 biological replicates). Statistical analysis was performed using Welch’s T-test. **(E)** Flow cytometry histograms of Citrine expression in WT and H3.3 K36M cells before KRAB recruitment (gray, -dox), after 6 days of dox silencing at 1000ng/ml dox (D6 +dox green, left), and 14 days memory after dox removal (20 day time point, right). Cells from these populations were collected for EMseq. **(F)** Top: Schematic of the AAVS1 reporter indicating regions for which DNA methylation is measured using EMseq. Bottom: **%** CpG methylation from EMseq plotted along the amplicon corresponding to the promoter region on the reporter upon KRAB recruitment and release at time points corresponding to populations shown in 4H (n=2 biological replicates). Shading represents the standard deviation. **(G)** Scatter plot measuring the time evolution of the mean %CpG methylation (n=2 replicates) at the promoter from EMseq (from Fig. 4H) and the mean % Citrine OFF from flow cytometry experiments (n=3 replicates taken from Fig. 1F) over time during KRAB silencing and memory. Y-axis error bars represent the SEM of 3 biological replicates. **(H)** Kinetic model describing steps associated with establishment and maintenance of epigenetic memory in parental and K36M cells In the first step (Establishment), H3K9me3 is converted to DNA methylation. H3.3 K36M cells (bottom) have reduced H3K36me3 and a reduced ability to convert H3K9me3 into DNA methylation (thinner arrow). (I) Schematic describing a possible model consistent with loss of epigenetic memory due to H3.3 K36M loss of H3K36 methylation due to K36M (right) leads to a reduced ability to recruit DNMT3A/B and hence low CpG DNA methylation.

To test whether levels of H3K36 methylation would impact establishment of epigenetic memory, we recruited KRAB for 2 days to silence the reporter, transfected the siRNAs targeting either NSD1 or SETD2 upon removal of dox, and measured reporter reactivation and memory over the course of 2 weeks with flow cytometry. Upon knockdown of either NSD1 or SETD2, more cells reactivated the reporter compared to cells treated with the scramble siRNA control **(Fig. 4C).** This phenotype is in line with the reduction of memory in cells harboring the K36M mutation. The steady-state memory in both NSD1 and SETD2 knockdown cells was consistently around 15% lower than the control cells, suggesting that H3K36 methylation is important for maintenance of KRAB mediated epigenetic memory–either directly or through co-recruitment of factors important for memory.

Given global H3K36 methylation loss was associated with gene reactivation after KRAB silencing at our reporter, we hypothesized that recruitment of protein co-factors responsible for establishment of epigenetic memory reliant on this modification was likely disrupted. H3K36me2 and H3K36me3 were both previously shown to bind to the PWWP domain of DNMT3A and DNMT3B, serving as a recruitment platform for de-novo DNA methylation, which is important for gene silencing and epigenetic memory.^20,71,72^. Furthermore, CpG methylation is important for robust and mitotically stable gene silencing in our reporter (**Fig. S4C**) and in most cell types. ^1,3,50,73–75^ Since KRAB-mediated epigenetic memory is dependent on DNA methylation ^51,75,76^, and since we observed a reduction of H3K36me3 by either expression of oncohistone or siRNA against NSD1 and SETD2, we hypothesized K36M could be disrupting the recruitment of DNMT3A or DNMT3B via their PWWP domains. This would eliminate the writing of DNA methylation needed to “lock in” the silenced state after KRAB recruitment. To test this hypothesis, we used siRNA against DNMT3A and DNMT3B individually or in combination after KRAB silencing to inhibit de novo methylation in the parental or H3K36M HEK293T cells **(Fig. 4D left, Fig. S4D)**. We reasoned that if recruitment of DNMTs is already inhibited in the K36M cell line, then further perturbation by siRNA would have minimal effects on epigenetic memory given our hypothesis that the oncohistone expression lowers the probability of DNMT3A/B recruitment at the reporter. However, we would expect that the siRNA would have a strong effect on the parental cell line which we hypothesize relies on DNMT3A/B for establishment of epigenetic memory. Knockdown of DNMT3A and DNMT3B after KRAB recruitment reduced epigenetic memory in the parental line, while the same knockdown had little effect on the H3K36 cells that instead maintained most of their starting memory (close to 5% difference from control). **(Fig. 4D** right). This was true for both individual siRNA’s against DNMT3A or DNMT3B **(Fig. S4E)**.

We next asked whether differences in response to DNMT3A/B knockdown are consistent with the H3.3 K36M cells having lower levels of CpG methylation at the reporter after KRAB recruitment. To measure methylation, we extracted genomic DNA at the end of silencing (at day 6) and after release once memory establishment is achieved (at day 20). We performed EM-seq (enzymatic conversion of unmethylated Cs to Us ^77^) followed by PCR amplification and sequencing of promoter and gene body regions in our reporter (**Fig. 4E)**. At the end of silencing, both the parent and K36M cell lines show very little DNA methylation at the promoter (**Fig. 4F middle**) or the gene body (**Fig. S4F**), comparable to the negative no-recruitment control (**Fig. 4F top**). However, by the day 20 memory timepoint, substantial methylation is present at the promoter (**Fig. 4F bottom**), suggesting that most de novo DNA methylation appears after removal of dox, during the memory phase. This is consistent with previous work in a different system, suggesting DNA methylation accumulates after continuous KRAB recruitment on the time-scale of weeks compared to gene silencing occurring within several days.^76^

We measured nearly twice the level of CpG methylation in the parental line compared to the K36M line at the promoter **(Fig. 4G),** indicating that accumulation of DNA methylation is blunted by the presence of H3.3 K36M and leads to a reduced number of cells with epigenetic memory. However, when normalizing the level of methylation to the fraction of silenced cells in the population used as input, the H3.3 K36M and Parent cells had similar levels of DNA methylation per cell. **(Fig. S4G)** This suggests that H3.3 K36M inhibits the fraction of cells that convert H3K9me3 to DNA methylation, but that once DNA methylation is acquired, the oncohistone no longer affects epigenetic silencing (**Fig. 4H, top**). From these data, we propose a model where H3K36me2/3 combines with H3K9me3 to recruit DNMT3A/B and establish DNA methylation at the promoter **(Fig. 4H, bottom).** Loss of H3K36 methylation due to oncohistone incorporation disrupts recruitment of DNMT3A/B, leading to fewer cells with DNA methylation, and hence reduced epigenetic memory after KRAB silencing.

### A chromatin spreading model that includes histone and DNA methylation quantitatively predicts decreases in epigenetic memory caused by H3.3 K36M

We next proceeded to test if the memory dynamics we observe in both the WT and oncohistone cells can be quantitatively explained by a stochastic model of chromatin modifications at an array of nucleosomes across the reporter. We used the basic framework of a previously established model used to explain the emergence of inherently bounded domains and the shape and size of steady-state H3K9 methylation profiles in MEFs and mESCs. ^78,79^ We modified this model ^79^ to incorporate a state corresponding to irreversible epigenetic memory, which we associate with DNA methylation, resulting in three possible states for each nucleosome in the array: active (A), reversibly repressed by histone modifications (R) and irreversibly repressed (I) (**Fig. 5A**). In our model, transitions between nucleosomal states occur in the form of four different reactions representing molecular actions of recruited KRAB and endogenous chromatin modifying enzymes on nucleosomes: 1) **Nucleation:** conversion from A to R for the nucleosome that is closest to the TetO binding sites (**Fig. 5A** blue arrows, **5B1**). This reaction only takes place in the presence of dox, i.e. KRAB recruitment of the TetO sites. 2) **Spreading:** local spreading of histone modifications where one randomly selected nucleosome in the R state recruits enzymes that attempt to modify one of the two neighboring nucleosomes (convert it from A to R) (**Fig. 5A** dashed arrow, **5B2**). Any one of the two immediately neighboring nucleosomes is chosen with equal probability for this conversion attempt. The spreading can happen in both dox and no dox conditions at the same rate. This is equivalent to spreading of both methylation and deacetylation. 3) **Erasure of repressive modifications:** direct R to A conversion, where one randomly selected nucleosome is attempted to be converted to the A state at a certain rate (which is only successful if the chosen nucleosome is in the R-state) (**Fig. 5A&B3**). This erasure is not dox dependent, but rather an intrinsic property associated with the promoter and active locus. 4) **Irreversible silencing:** direct R to I transition where a randomly selected nucleosome at the promoter (here consisting of 5 nucleosomes) is attempted to be converted to an I state (**Fig. 5A&B4**). We chose to allow the irreversible conversions only at the promoter because we measure changes in methylation primarily in this region (**Fig. 4G&S4F**).

**Fig. 5.**
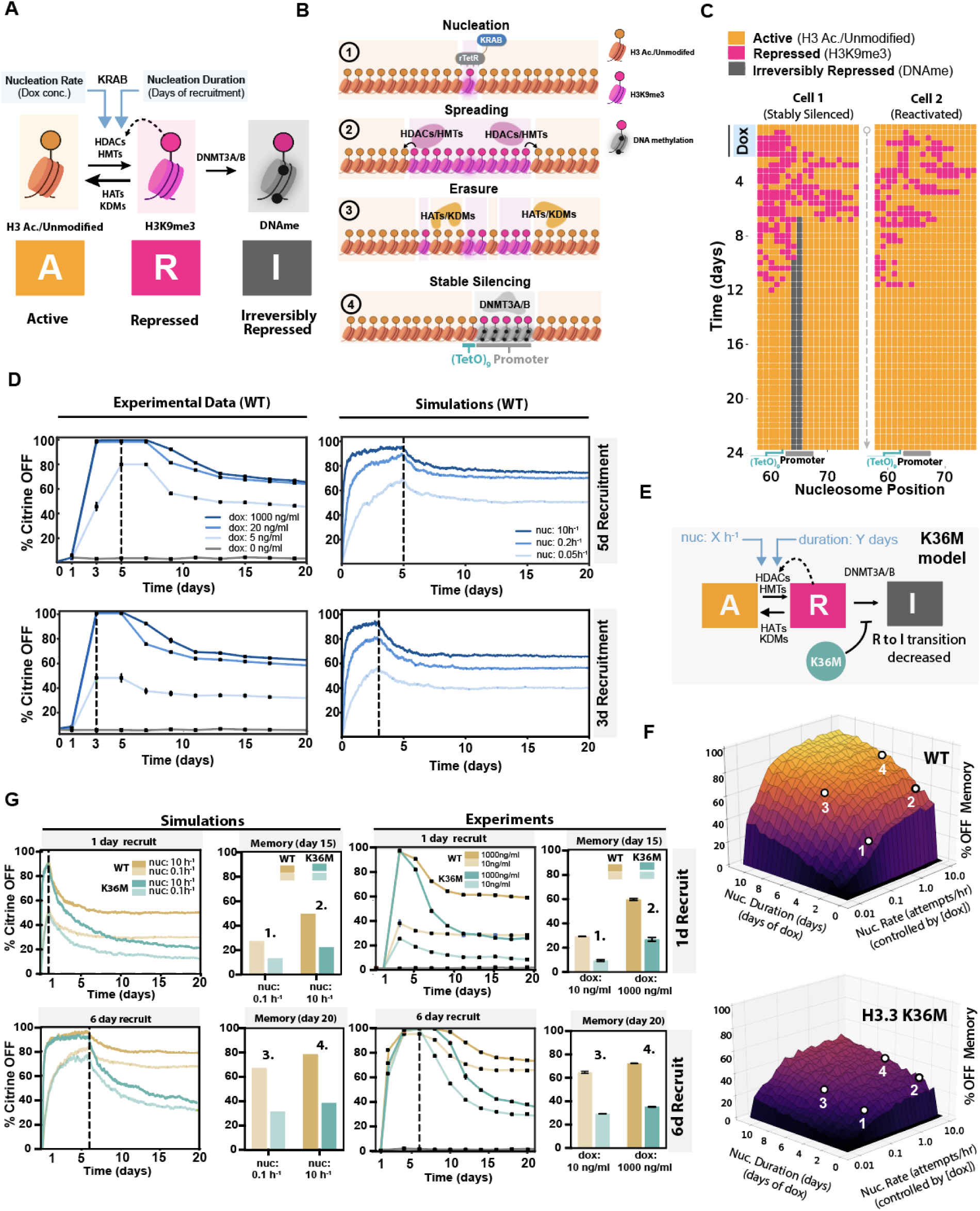
A chromatin spreading model that includes DNA methylation quantitatively predicts decreases in epigenetic memory caused by H3.3 K36M. **(A)** Schematic showing three nucleosome states: Active modifications (A, yellow, e.g., H3 acetylated/H3K9 unmodified), Repressive modifications (R, pink, e.g., H3K9me) and Irreversibly Repressive modifications (I, gray, e.g. DNA methylation). Example enzymes that mediate state conversions between nucleosome states are indicated on the black arrows that denote state transitions among the 3 states. KRAB recruitment increases the rates of transition from A to R (at the nucleosome where KRAB is recruited), indicated by the blue arrow. We call this KRAB-mediated increase in A-to-R conversion nucleation; dox concentration and recruitment time increase nucleation rate and duration respectively. **(B)** Typical time-evolution of the nucleosome array: 1) The system starts with all nucleosomes in the active state (yellow) and upon dox addition KRAB binds at the nucleosome corresponding to the 9XTetO sites to nucleate H3K9me3 (R state conversion, pink). 2) H3K9me3 (R state, pink) spreads from the nucleation site through reader-writer feedback between neighboring nucleosomes away from the nucleation site. 3) Spreading of repressing modifications can be counteracted (“erased”) by direct R to A nucleosome state transitions coming from histone turnover, removal of histone methylation, or deposition of histone acetylation. 4) Nucleosomes at the promoter can be irreversibly silenced (gray) by acquiring DNA methylation. **(C)** Stable repression is stochastic in individual simulations under the same conditions (rates). Left: Example of one cell with an irreversibly silenced reporter gene due to the acquisition of nucleosomes in I state (gray) at the promoter. Right: Example of another cell whose reporter becomes reactivated (all nucleosomes in the yellow state) after transient silencing. **(D)** Left: Experimental time-series data showing the fraction of silent cells measured every other day from the time point at which dox has been added for a transient period - either for a total of 5 days (top panel) or 3 days (bottom panel), up to dashed line- and at different concentrations (in ng/mL), as shown in legend of lower panel (n=3 biological replicates). Dox was removed for days to the right of the dashed line. Right: Simulation data showing the fraction of silent reporters (from 1000 simulations) every 20 minutes (in simulation time) for different nucleation rates that were kept constant for either 5 days (top) or 3 days (bottom). Nucleation rates were set to zero for times to the right of the dashed line. **(E)** Schematic showing the kinetic model used for generating the memory landscapes of parental (WT) cells and oncohistone cells in F. Input variables are the nucleation rate (varied on a log-scale from 0 ℎ^−1^ to 10 ℎ^−1^) and nucleation days (varied from 0 to 11, in steps of 0.5 days). The oncohistone simulations were performed with the same parameters but with 75% of randomly chosen nucleosomes being blocked for R to I transitions to mimic the rate-reducing effect of the oncohistone on de novo DNA methylation. **(F)** Memory-landscapes of reporter gene in parental cells (top) and H3.3K36M oncohistone cells (bottom) showing the fraction of silent cells at simulation day 14 as a function of nucleation-rate (in attempts/hour) and nucleation duration (in days). **(G)** Left: Simulation time courses performed with nucleation rates of 0.1 ℎ^−1^(transparent) and 10 ℎ^−1^(nontransparent) and with 1 (top) and 6 days (bottom) of nucleation for oncohistones (blue) and WT cells (mustard), respectively. Barplots show the fraction of silenced cells at memory day 14. Right: Experimental time courses performed with 10 ng/mL (transparent) and 1000 ng/mL (nontransparent) of dox for 1 (top) and 6 days (bottom) for oncohistones (blue) and parental WT cells (mustard), respectively. Barplots show the fraction of silenced cells at memory day 14.

We simulate the dynamics of nucleosome modifications in each cell using a one-dimensional 128 nucleosome array (which is sufficiently long to exclude any boundary effects) with a centrally placed nucleation site (rTetR-KRAB recruitment) and adjacent promoter. At each step, one of the four reactions is selected as an event using the Gillespie algorithm^80^, and nucleosomes are subsequently chosen randomly for a nucleosome-state conversion attempt as described above **(see methods for details)**. Two representative examples of a simulated time-evolution of the system illustrate the behavior of the model (**Fig. 5C**, zoomed in on the 20 central nucleosomes). At the beginning (top), all nucleosomes are in the A state; upon nucleation, R nucleosomes start spreading from the nucleation site while being counteracted by erasure (direct R to A transitions). Depending on the length and the rate of nucleation, some of the R nucleosomes at the promoter are irreversibly converted to I nucleosomes stochastically in some simulations (**Fig. 5C, left**), while in other simulations the array does not acquire any I nucleosomes and revert back to the active state (**Fig. 5C, right**).

We first manually fitted parameter values for the spreading rate, the erasure rate, and the stable silencing rate at each nucleosomal position such that the simulated data matches the WT experimental data for different concentrations and durations of dox recruitment (**Fig. 5D**, left). To mimic the experiments, the nucleation rate (corresponding to dox concentration) and its duration (recruitment time) were the only parameters allowed to change between the different curves/experiments while all other parameters were fixed. We assumed that the nucleation rate (in ℎ^−1^) is directly proportional to the dox concentration (see Methods). With these assumptions, we were able to find model rates that recapitulated the experimental results of both silencing and memory we measure for different dox concentrations and durations of recruitment (**Fig. 5D**, right.)

Our model requires two major assumptions: 1) The spreading rate must be precisely balanced with the erasure rate to preserve a maximal silencing potential of the promoter while ensuring effective erasure of histone methylation after the nucleation phase. If the spreading rate is too high compared to the erasure rate, we find cells cannot reactivate, leading to 100% irreversible silenced memory; on the other hand, if turnover is increased compared to the spreading rate, cells are unable to be effectively silenced (**Fig. S5B-D**). Thus, the system exists very close to a critical point before loss of inherent confinement ^78,79^ **and Fig. S5A-D**). 2) The promoter must be very sensitive to R and I nucleosomes with one R or I nucleosome being sufficient to silence the whole promoter. If we require gene silencing to necessitate more nucleosomes in the repressed states (R or I), full promoter silencing is not achievable for rates that allow reactivation (**Fig. S5E&F**).

Next, we modified the model to account for the presence of the H3.3 K36M oncohistone. This was based on existing literature and our data that suggest H3K36me2/3 and H3K9me3 work in tandem to recruit de novo DNA methylation. We simulated oncohistone incorporation by randomly replacing 75% of the wildtype nucleosomes on the array with H3.3K36M-containing nucleosomes that are unable to transition from R to I state. (**Fig. 5E)**. This blockade of nucleosomes preventing transition to irreversible silencing state effectively mimics an impaired rate of de novo DNA methylation due to a loss of H3K36me2/3. The other rates in the model were kept the same for the H3K36M as in the WT. Using this approach, we generated two epigenetic memory landscapes, in WT and H3K36M cells, that make predictions about the epigenetic memory (fraction of cells with the reporter silenced 14 days after KRAB recruitment/nucleation) over a large parameter space of KRAB recruitment duration and dox concentration.(**Fig. 5F, Fig. S5H**). While the WT memory landscape predicts a high fraction of permanently silenced cells for longer nucleation durations and rates (>60%), the H3.3K36M cells never reach more than 50% silenced cells, even for long nucleation durations and high nucleation rates (10 ℎ^−1^). Therefore, the H3K36M cells are predicted to show a systematic reduction of epigenetic memory across conditions.

To experimentally validate predictions made in the epigenetic memory landscapes, we manually picked 4 points from the plot surfaces (indicated as points 1 through 4 in **Fig. 5F**) representing different nucleation strengths and durations of KRAB recruitment and a sufficient dynamic range in memory to be resolved using flow cytometry. Given nucleation strength and nucleation duration are reflective of different concentrations of doxycycline and the duration of dox applied to cells respectively, we were able to map the parameter space onto experimental conditions predicted by the simulations (see methods for details). For these 4 sets of parameters (see number labels on plots corresponding to conditions selected from **Fig. 5F**), the predicted simulation time courses for oncohistone and parent cells (**Fig. 5F left**) are in close agreement with experiments for both silencing and memory dynamics (**Fig. 5F right)** suggesting the model robustly predicts epigenetic memory over a wide parameter space of recruitment strength and duration.

## Discussion

In this study we show that H3.3 K36M expression and the associated decrease in H3K36me2/3 diminish epigenetic memory and increase reactivation after KRAB mediated gene silencing in both HEK293T cells and a TC28a2 chondroblastoma model. This finding highlights a new role for H3K36 methylation in epigenetic memory and illustrates the versatility of H3K36 methylation states in mammals– specifically its ability to not only be involved in active transcription, including splicing and transcriptional elongation, but also to serve functions involved with gene repression. We additionally show that the mutant H3.3 K36M histone diminishes memory formation through inhibition of DNA methylation. Moreover, we can accurately predict KRAB-mediated silencing and memory in both the WT and K36M using a stochastic model that simulates spreading of histone methylation across the locus, and impaired conversion of this repressive state to an irreversible silenced state (associated DNA methylation) in the oncohistone presence.

When H3.3 K36M is introduced through endogenous mutation or exogenous overexpression, we observe both H3K36me2 and H3K36me3 are reduced globally in both HEK293T and TC28a2 cells, due to inhibition of H3K36 methyltransferases SETD2 and NSD1/2/3 by the oncohistone. ^10,11,60^ Despite the aforementioned changes in histone modifications, we note that differences in epigenetic memory did not depend on H3K9me3, as the observed levels of H3K9me3 upon KRAB recruitment and release were identical between WT and H3.3 K36M cells. However, we find that promoter DNA methylation after KRAB recruitment and memory establishment (KRAB release) was lower in the H3.3 K36M cells compared to the parental line. Given that H3K9me3 mediated heterochromatin has been shown to lead to DNA methylase recruitment ^40,41^, and our data shows differences in epigenetic memory inhibition between H3.3 K36M and WT cells upon siRNA against DNMT3A/B, we hypothesize that the oncohistone likely blocks the ability of DNMT3A/B to methylate DNA through reduced recruitment of these methyltransferases. This is in line with previous findings that showed the PWWP domains of DNMT3A and DNMT3B, are important for effective binding of DNA through recognition of H3K36me2/3 methylation. ^20,71,72^

Despite the fact that H3K36 methylation is commonly associated with regions of active transcription, our work in mammalian cells reinforces findings from previous studies in drosophila and yeast where H3K36 methylation has been shown in several contexts to have the ability to function with heterochromatin to mediate transcriptional repression. ^11,14,18,19^ We show H3K36me2/3 can act in tandem with H3K9me3 to deposit DNA methylation and maintain stable and robust maintenance of gene silencing. Although the expression of H3.3 K36M has been shown to cause global increases of PRC2-mediated H3K27me3, a repressive histone modification also involved in heritable gene silencing, we exclude this as a possible cause of silencing or memory changes at our reporter, since we did not detect any significant changes upon dox mediated KRAB recruitment in either cell line **(Fig. S3F,G).** Furthermore, increases of H3K27me3 would be predicted to result in an increase in epigenetic memory in the oncohistone lines, which is contrary to our observations of reduced memory. Thus, we propose the disruption of DNA methylation deposition, through H3K36me2/3, as the likely mechanism by which the K36M oncohistone causes the decrease in memory establishment.

We note that despite the fact that KRAB leads to silencing in nearly 100% of the cells both after 2 days and 6 days of recruitment, most DNA methylation observed at the promoter accumulates after KRAB release, between Day 6 and Day 20 **(Fig. 4F)**. These timescales of DNA methylation deposition are consistent with previous observations of similarly slow build up of CpG methylation on the time-scale of weeks^76^; however, in those studies KRAB was continuously recruited for weeks, in contrast to our work where we transiently recruit for days and then follow memory after release. Our observations of DNA accumulation after KRAB release suggest that molecular interactions involved in DNA methylation persist well after dox is removed and rTetR-KRAB is no longer bound to the reporter. Given that the average promoter DNA methylation when normalized to the fraction of cells that are irreversibly silenced is nearly identical between H3.3 K36M and the parental line, it suggests that cells that are irreversibly silenced have accumulated similar amounts of methylation regardless of oncohistone expression. It is likely that H3.3 K36M thus controls the fraction of cells that convert histone methylation to DNA methylation, but that once DNA methylation is acquired in a cell, it leads to permanent memory in both the WT and oncohistone lines (**Fig. 4H**).

Colocalization of H3K9me3 and H3K36me3 has been previously reported in mammalian cells in regions coined “atypical heterochromatin” ^17^, and this colocalization was dependent on both the SETDB1 (H3K9me3 writer) and NSD enzymes (H3K36me3 writers). These regions were previously characterized as precursors to enhancers which would activate once SETDB1 was removed. Our study adds to our understanding of a similar type of heterochromatin and how H3K9me3 and H3K36 methylation can act in tandem to establish memory of gene repression via recruitment of DNA methylation if these domains last for long enough. This slow conversion of H3K9me3 at previously transcribed genes (containing H3K36 methylation) to DNA methylation could serve as a timer that allows gene reactivation within a certain period, but can still allow commitment to irreversible fate transitions during cell differentiation for example.

Recent studies have confirmed that H3K36 methylation plays an important role in maintenance of cell identity during development and differentiation, not only by blocking the repressive effects of PRC2 at lineage-specific genes, but also by repressing alternative lineage-specifying genes through exclusion of transcription factors in a DNA-methylation-dependent manner.^81^ This observation is consistent with our finding that the interplay between DNA methylation and H3K36 methylation tunes the rate of irreversible fate transitions.

Given that often multiple histone modifications occur simultaneously to achieve their function at endogenous genes (e.g., bivalent chromatin), we anticipate that our finding that H3K36 methylation contributes to epigenetic memory establishment will aid in the development of epigenetic tools. For example, in order to increase epigenetic memory after transient targeting of endogenous genes using CRISPR (dCas9), recent work combined KRAB with DNMT3A and DNMT3L.^73,82^ Recent work has already shown that dCas9 fused to SETD2 by itself was sufficient to repress gene expression.^83^ Based on these findings and our data, it seems likely that combining the catalytic domains of H3K36 methyltransferases to commonly utilized repressive domains associated with H3K9me3, such as KRAB domains, may result in more robust and stable epigenetic memory after gene silencing at endogenous regions. Thus, future potential combinations of H3K36 methyltransferase domains with transcriptional repressors may improve upon current standard epigenome targeting strategies in locations that are difficult to silence, and may ensure more robust epigenetic memory.

In general, losing the ability to reliably establish or maintain epigenetic memory, and hence cell identity, through cell divisions can be a driver of cancers. ^84^ In a recent study, cells carrying mutations in MCM2, a protein associated with recycling of histones after DNA replication in eukaryotes, led to defects in differentiation and tumor progression in cell culture models due to impaired histone inheritance.^85,86^ While here we used our recruitment system to study establishment and maintenance of epigenetic memory in the context of a particular cancer-causing histone mutation, our approach can be extended to study this phenomenon in other cancer types.

## Methods

### Cell Culture

HEK293T cells and TC28a2 cells were cultured with DMEM GlutaMAX (Gibco, 10-566-024) supplemented with 10 % TET approved FBS (Omega Scientific FB-15 or Sigma Aldrich F0926) and 1 % Penicillin-Streptomycin (Gibco, 15-140-122). Cells were kept in a humidified incubator at 37°C and 5 % CO2. H3.3 K36M or cells overexpressing histones were continuously grown in the presence of 50μM Indole-3-acetic acid (IAA) (Sigma-Aldrich, I5148-2G) until experiments were performed to suppress protein expression. All cell lines routinely tested negative for mycoplasma.

### Lentivirus Production and Transduction

Lentivirus was generated in a 10cm dish format by transfecting HEK293T Lenti-X cells (Takara) at roughly 80% confluency with 4.5μg of an equimolar mixture of three plasmids encoding the envelope and viral packaging components from the laboratory of Didier Trono (pMD2.G: Addgene 12259; pRSV-Rev: Addgene 12253; pMDLg/pRRE: Addgene 12251). 4.5μg of lentiviral donor plasmid was added to the mixture and incubated for 20 min with 10μL of polyethylenimine (PEI, Polysciences 23966) dissolved in OptiMEM (Gibco 31985062). Lentivirus-containing cell culture supernatant was harvested 72 hours post transfection. Harvested virus was 0.4μM filtered (Celltreat 229749) and concentrated by centrifugation using a 100kdA cut-off PES protein concentrator (Pierce 88533) prior to storage at -80C. Lentivirus was added dropwise to cells during transduction and incubated for 24 hours prior to wash off and propagation of cells.

### Cell Line Generation

HEK293T reporter lines were generated using TALEN-mediated homology-directed repair to integrate donor constructs into the AAVS1 locus by reverse transfection of 1E6 HEK293T cells with 5ul Lipofectamine LTX (Invitrogen 15338030) supplemented with 2.5ul PLUS reagent with 1000 ng of reporter donor plasmid and 500 ng of each TALEN-L (Addgene 35431) and TALEN-R (Addgene 35432) plasmid targeting upstream and downstream the intended DNA cleavage site. Cells were treated with 1000 ng/mL puromycin (Invivogen ant-pr-1) for 5-7 days to select for a population where the donor was stably integrated in the intended locus prior to FACs enriching Citrine positive cells. Reporter lines were then transduced with lentivirus containing osTIR1-9xMyc which was subcloned from pBabe Puro osTIR1-9xMyc from the laboratory of Andrew Holland (Addgene 80074) and selected with 7.5 μg/mL Zeocin (Invivogen ant-zn-05) for 10 days. These cells were then transduced with lentivirus expressing H3.3 K36M or analogous histone vector and selected with 10μg/mL Blasticidin (Gibco A1113903) for 10 days prior to FACs enrichment based on mCherry signal. Recruiter plasmids rTetR-KRAB and rTetR-VP64 were introduced in a similar manner except cells were FACS sorted without selection 72hr post transduction for appropriate fluorescence signal on the vectors. TC28a2 reporter lines were generated using an identical manner to HEK293T except DNA was introduced using electroporation using the X-001 program on the Nucleofector 2b (Lonza, AAB-1001) in combination with Nucleofector Kit V (Lonza, VCA-1003) according to the manufacturer’s recommended protocol.

### Flow Cytometry

Flow cytometry during recruitment assays for activation or repression were performed in a 24-well format where 15,000-20,000 cells were seeded per well and grown in the presence of doxycycline hyclate (dox) between 5ng/ml to 1μg/ml (Tocris 4090). Dox was replenished daily when concentrations of less than 100 ng/ml were used to account for drug degradation. In all other instances, fresh media containing dox was replenished every 2 days with cell passaging. For memory timepoints, dox was washed out and cells were continued to be grown in the presence of DMEM growth medium. For cytometry measurements, cells were washed 1x with 200μl DPBS (Thermo Fisher 14190250) followed by incubation with 80μl 0.25% Trypsin (Thermo Fisher 252-000-56) for 5 minutes. Cell suspension was diluted with 200μl DMEM media. 150ul of the suspension was filtered through a 40μM cell strainer at 40G for 5 minutes to remove clumped cells and subsequently used for analysis on the ZE5 flow cytometer (Biorad). Analysis of gene expression was analyzed using the Easyflow MATLAB script (https://antebilab.github.io/easyflow/). To all data, singlet cells expressing the rTetR recruiters were first selected based on forward and side scatter gates and mTurquoise fluorescence. To these cells a tertiary gate was applied to oncohistone expressing cells when appropriate to select for mCherry positive cells. A Citrine OFF% was then calculated by manual gate being imposed on the FITC channel so that roughly no more 1-10% of the mCitine signal was positive in the untreated (no dox) population for all samples. In TC28a2 cell experiments, memory experiments were normalized to the no dox control to account for higher background silencing of the reporter. This was performed by calculating the percentage of OFF cells as follows at each timepoint: 100* (%OFF*no do*x - %OFF*dox)***/**(100-%OFF*no dox*)

### siRNA Transfection

siRNA transduction of HEK293T cells were performed in a 24-well format using Lipofectamine RNAiMax (Invitrogen 13778150) according to manufacturer’s protocols. Briefly, 15,000 cells were reverse-transfected with 0.3μl of Lipofectamine RNAiMax complex containing either 1 pico mol Silencer siRNA targeting or control non-targeting siRNA (Invitrogen 4390843). Media was exchanged 24 hours later and gene expression was assessed beginning 48 hours later in flow cytometry time courses or used as an endpoint for western blots. siRNA for western blot validations were performed similarly as above but scaled up to multiple wells of 6 well plate format according to manufacturer protocol where roughly 375,000 cells were transfected and 50 pico mol of siRNA per well and pooled together for analysis. The following siRNA were used in the study from the Invitrogen pre-designed siRNA series (cat# 4427036/4427037): siNSD1(s34629), siNSD2(s200460), siNSD3(s29725), siSETD2 (s26423), siDNMT3A (s200426), siDNMT3B (s4222).

### Western Blotting

5-20 million cells were lysed in ice cold RIPA buffer (Millipore Sigma 20-188) in the presence of protease inhibitor cocktail (Roche 11697498001). Cell lysis was carried out with nutation for 30 min at 4C followed by homogenization for 15 minutes on the Covaris E220 sonicator set to Peak Power: 140-175.0, Duty Factor: 10.0 and Cycles/Burst:200. Protein loading was equilibrated by measuring concentration on a Qubit using the Protein BR kit (Thermo Fisher Q33211). 20-30μg lysate was boiled for 10 minutes with loading dye (Thermo Fisher 26634) and run on 4-20% Tris/Glycine gels (Biorad 4568096). Transfer was performed using the Trans-blot turbo system (Biorad) with PVDF membranes (Biorad 1704156). Membrane was probed using primary antibodies at 1:1000 dissolved in blocking buffer (Thermo 37543) overnight at 4C with shaking. Secondary ECL HRP-conjugated antibodies (Cytiva NA934 or NA931) were used at 1:2000 and incubated for 1 hour at room temperature with shaking. Blots were developed with HRP substrate (Millipore Sigma WBKLS0500) and imaged on the iBright imaging system (Thermo Fisher FL1500). A Histone H3 (EMD Millipore 07-690) antibody was used as a loading control at 1:5000 dilution. When necessary, PVDF membranes were stripped using a stripping buffer (Thermo 46430) according to the manufacturers protocols and re-probed with antibody for a subsequent round of imaging. Target antibodies used are as follows: HA-tag (Cell Signaling 3724), H3K36me2 (Cell Signaling), H3K36me3 (Abcam ab9050).

### Magnetic Cell Separation

Cells were detached from tissue culture plates by washing once with DPBS (Gibco 14190-250) and incubating with TypLE Express (Gibco 12604021) for 5 minutes at 37C. Cells were then pelleted and washed 2x with DPBS to remove residual trypsin and medium. ProteinG dynabeads (Thermo Scientific 10003D) was used at a concentration of 10μl beads per 1E6 cells. Beads were washed twice with 5x volume of magnetic separation wash buffer (2% BSA in DPBS) on a magnetic stand. Washed beads were then incubated with resuspended cells for 90 minutes at room temperature with nutation in magnetic separation wash buffer at 10x volume of buffer to beads to allow binding of the IgG surface marker to beads. The solution containing cells and beads was then placed on a magnetic stand for 5 minutes to separate bound and unbound cells. The supernatant (unbound cells) was transferred to a new tube and re-applied to the magnet to remove residual beads, and the bound fraction was washed once more at 5x volume magnetic separation wash buffer to remove residual non-bound or non-specifically bound cells on the magnet and resuspended in 5x buffer. The enriched bead slurry containing bound cells and the unbound supernatant were sampled directly via flow cytometry to assess sorting efficiency and subsequently placed into separate tissue culture vessels and propagated using normal cell culture methods to assess epigenetic memory.

### DNA methylation sequencing

Genomic DNA was extracted from roughly 500,000 HEK293T cells using the Monarch Genomic DNA Purification Kit (NEB T3010L) and fragmented by digestion with XbaI and AccI to excise and release the genomic DNA region corresponding to the AAVS1 reporter. The DNA was then size selected using 1x volume SPRI bead clean up (Beckman Coulter B23317). 200ng of cleaned DNA was used as an input for conversion using a Enzymatic Methyl-seq Kit (NEB E7120S) according to the manufacturer’s recommended protocol. Briefly, APOBEC converted DNA was PCR amplified using 22x cycles with Q5U master mix (NEB M0597S) with primer pairs cTF378+cTF381 for the promoter and oSA021+oSA023 for the gene body. Amplified DNA was then size enriched using 0.9X volume SPRI beads and subject to another round of PCR using 10x cycles with NEBNext Ultra II (NEB M0544S) using primers pairs cTF383+cTF399 for the promoter and oSA021+oSA023 for the gene body to add Illumina Read1 and Read2 overhangs. DNA was cleaned up using 0.9X volume SPRI beads. The above PCR and clean up steps were repeated for a third PCR reaction with indexing primers to append Illumina indices for sequencing. Pooled and indexed DNA was sequenced using a MiSeq 600 cycle kit (Illumina MS-102-3003) reading 374 cycles in read1 and 235 cycles in read2. BWAmeth was used for methylation-specific alignment to the reporter, and samtools was used to convert raw alignment files to indexed .bam. Bulk CpG methylation levels were calculated per site using MethylDackel (https://github.com/dpryan79/MethylDackel). Custom analyses were used (dSMF-footprints_optional_clustering.py) to produce single-molecule methylation heatmaps of aligned reporter molecules. Subsequent analysis and visualization was performed in Python.

### CUT&RUN

CUT&RUN was performed with the CUTANA ChIC/CUT&RUN kit (EpiCypher 14-1048) according to manufacturer’s instructions from manual version 3, with modifications as follows. 500,000 HEK293T cells were thawed and used directly from -80C as input for all reactions. Cells were permeabilized using 0.025% digitonin. Antibody incubations were all performed overnight at 4C (see antibody list) and used at 1:50 (1μl per reaction) with the exception of H3K9me3 which was used at 1:33 (1.5μl per reaction). Libraries were prepared using the CUT&RUN Library Prep Kit (EpiCypher 14-1001/14-1002) with 1-20ng of input DNA according to manufacturer’s protocols. DNA was quantified using a Qubit4 fluorometer and libraries were sequenced at a targeted depth of 10M-20M paired end reads per sample on either an Illumina NextSeq500 (High Output, 2×36 Cycles) or HiSeq2500 (2×150). Normalization of sample inputs was performed when applicable either using Spike-in (either from reads from E Coli genome or barcode reads from the K-MetStat panel (EpiCypher 19-1002)) in accordance with manufacturer’s instructions or CPM normalization to total reads when applicable. Antibodies used for CUT&RUN are as follows: H3K9me3 (Abcam ab176916), H3K4me3 (EpiCypher 13-0041), H3K27me3 (Cell Signaling 9733), HA-tag (Cell Signaling 3724).

### CUT & RUN data analysis

CUT&RUN sequencing data were aligned, filtered, and de-duplicated as described in a previous study ^28^. Specifically, a custom genome with our reporter sequence appended to the end of the hg19 human genome assembly was constructed with bowtie2-build. Paired-end alignment was performed with the following bowtie2 command: bowtie2 --local --very-sensitive-local --no-unal --no-mixed --no-discordant --phred33 -I 10 X 700 × {reference genome} -p 8 – 1 {first mate of pair} – 2 {second mate of pair} -S {output SAM file name}. Fragments that mapped completely within non-unique reporter elements (i.e. pEF, AmpR, or SV40 polyA) were ambiguous and thus removed to avoid confounding. Samtools was used to convert SAM files to BAM files and to subsequently sort and index BAM files. Picard was used to mark and remove duplicates with the following command: java -jar {picard tool} MarkDuplicates -I {input sorted BAM file} -O {output deduplicated BAM file} -M {output metrics file} --REMOVE_DUPLICATES true.

Data were normalized by one of three methods, as indicated in the text: counts per million (CPM), *E. coli* DNA spike-in, or K-MetStat panel spike-in. CPM normalization accounts for differences in sequencing read depth, required no changes to the experimental workflow, and used the following bamCoverage command: bamCoverage --bam {input deduplicated BAM file} -o {output bedgraph file} -- outFileFormat bedgraph --extendReads --centerReads --binSize 10 --normalizeUsing CPM. *E. coli* DNA spike-in normalization required adding the recommended amount of *E. coli* Spike-In DNA (provided with the EpiCypher kit) into the STOP buffer for quenching pAG-MNase activity. Because the same amount of *E. coli* DNA is added to each sample, this normalization method accounts for differences in sample processing from genomic DNA extraction onward. Scaling factors for normalization were calculated as the inverse of the percentage of uniquely aligned reads in each sample that mapped to the *E. coli* K12 MG1655 reference genome, as defined in the EpiCypher manual, and normalization used the following bamCoverage command: bamCoverage --bam {input deduplicated BAM file} -o {output bedgraph file} -- outFileFormat bedgraph --extendReads --centerReads --binSize 10 --scaleFactor {*E. coli* spike-in DNA scale factor}. K-MetStat panel spike-in normalization required adding a panel of DNA-barcoded, post-translationally modified nucleosomes conjugated to magnetic beads to each cell sample prior to antibody addition. This panel can provide information regarding specificity of antibodies against common histone modifications and can be used for normalization within an antibody condition. Histone modification-specific DNA barcodes were quantified for each sample using the shell script at epicypher.com/14-1048. Scaling factors for normalization were calculated as the inverse of the percentage of reads in each sample containing DNA barcodes for the expected histone modification, and normalization used the following bamCoverage command: bamCoverage --bam {input deduplicated BAM file} -o {output bedgraph file} -- outFileFormat bedgraph --extendReads --centerReads --binSize 10 --scaleFactor {K-MetStat scale factor}. Peak-calling was performed with SEACR ^87^ using the following command: bash SEACR_1.3.sh {experimental bedgraph} 0.01 non stringent {output prefix}. Peaks were annotated with the nearest gene and associated genomic feature using the R Bioconductor package ChIPseeker ^88^. CUT&RUN data were visualized via the Broad Institute and UC San Diego Integrative Genomics Viewer. Additional data analyses and visualization were performed with custom scripts in Python.

### RT-qPCR

RT-qPCR for siRNA validation was performed by extraction of RNA using the RNeasy Mini kit (Qiagen 74104) according to the manufacturer’s recommendations 24hr post siRNA transfection. cDNA synthesis was performed using iScript reverse transcriptase master mix (Biorad 1708840). qPCR was performed using SSO Advanced SYBR Green master mix (Biorad 1725271) on a CFX Connect Real-Time PCR system (Biorad 1855201). qPCR primers used for DNMT3A and DNMT3B respectively are as follows: **D3A FWD#2** 5’ TCTGGAGCATGGCAGGATAG 3’ **D3A REV#2:** 5’ AAATGCTGGTCTTTGCCCTG 3’ **D3B FWD#1** 5’ TTGCTGTTGGAACCGTGAAG 3’ **D3B REV#1** 5’ CCGCCAATCACCAAGTCAAA 3’

### Cell growth assays

Cell growth quantification was performed using the ViaFluor 488 SE cell proliferation dye in minCMV reporter HEK293T cells (Biotium 30086) or CellTrace Far-Red proliferation dye in pEF reporter TC28a2 cells (Thermo C34564) according to manufacturer’s instructions. Briefly, approximately 1E6 cells were stained for 20 min at 37C and centrifuged and washed two times with DMEM media prior to being monitored by flow cytometry over the course of several days of proliferation to measure fluorescence intensity due to cell division.

### Spreading Model Assumptions and Rates

KRAB leads to recruitment of HDACs and HMTs (SETDB1) leading to removal of active modifications (A, eg. histone acetylation) and deposition of repressive ones (R, e.g., H3K9 methylation) at the nucleosome closest to the TetO sites. ^46,89^ For simplicity, deacetylation and H3K9 methylation are absorbed into one transition (A to R conversions) where H3 deacetylation and H3K9me marks then spread via a read-write mechanism where chromatin modifiers bind to histones that are modified with the same substrate that they themself then deposit on neighboring nucleosomes. ^90,91^ Here, we assumed spreading of modifications happens linearly, meaning only between neighboring nucleosomes. It has been reported in the literature, that at certain genomic regions, DNA methylation occurs slowly and in an H3K9me dependent manner at promoters that are thought to reinforce the permanent silencing/heterochromatinization of genes. ^40^ At each nucleosome, the conversion from the repressed histone modifications state R to the irreversible state I is independent of KRAB recruitment. H3K9me is counteracted by HDMs, HATs and histone turnover. These biological observations formed the basis of our model, which is based on the following assumptions:

1. The system of 128 nucleosome positions contains two special regions - the centrally placed nucleation site at position 62 and the adjacent promoter region comprising nucleosome positions 63 to 67 which determines the state of the Citrine promoter.
2. The Citrine reporter is marked as silent when one or more nucleosomes in the promoter regions are in either the R or the I state.
3. Nucleation (direct A to R transitions) only occurs at the nucleating nucleosome (number 62). All other nucleosome positions have a nucleation rate of 0.
4. The average conversion rate between R and A nucleosomes is 0.33 attempts per hour at every nucleosome in the system, regardless of its position.
5. Spreading (positive feedback) reactions, specifically R-mediated A to R conversions, only take place between adjacent nucleosomes. The average attempt rate for these reactions is 1 per hour per nucleosome, regardless of the nucleosome’s position.
6. I nucleosomes are irreversible meaning that they do not change their state spontaneously.
7. Stable silencing (direct R to I transitions) only occurs at the five promoter nucleosomes at a low rate of 0.0025 attempts per hour at each position (if not blocked as in the oncohistone simulations).
8. Only R nucleosomes can spread through local R-mediated A to R conversions. A and R nucleosomes do not mediate local positive feedback reactions.

**Table S1.**
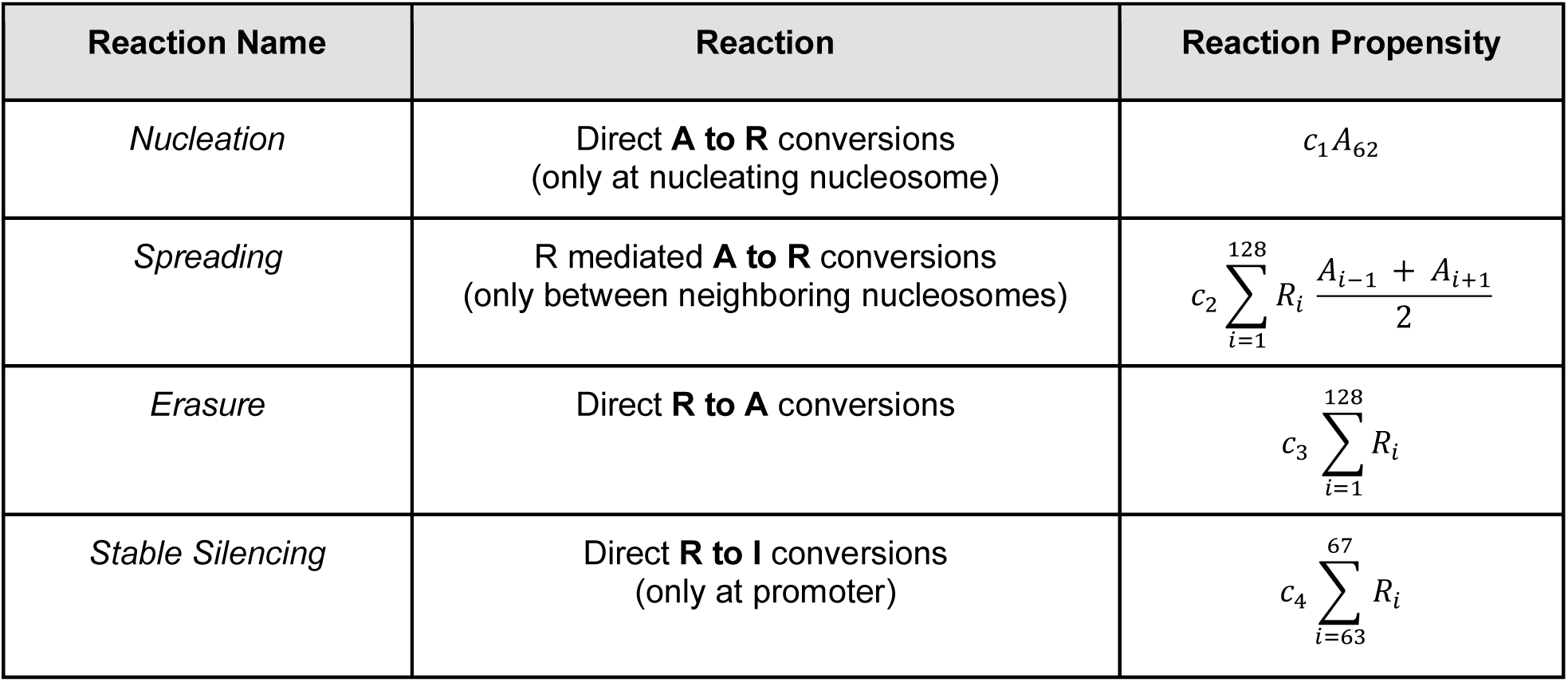
Rates used in the spreading model. A, R and I are the 3 nucleosome states shown in Figure 5A*. A_i_* and *R_i_* indicate that the nucleosome at position *i* in the nucleosome array is in state A or R respectively. *A_i_* is 1 if the nucleosome at position *i* is in the active state and zero otherwise. *A_62_* denotes the nucleosome at the site of nucleation where KRAB is recruited. *c*_1_ is allowed to vary between 0.05 ℎ^−1^ and 10 ℎ^−1^ (to mimic dox concentration changes), *c*_2_ is fixed at 1 ℎ^−1^, *c*_2_ is fixed at 1 ℎ^−1^, *c*_3_ is fixed at 0.33 ℎ^−1^ and *c*_4_ is fixed at 0.0025 ℎ^−1^.

| Reaction Name | Reaction | Reaction Propensity |
| --- | --- | --- |
| *Nucleation* | Direct **A to R** conversions<br>(only at nucleating nucleosome) | $c_1 A_{62}$ |
| *Spreading* | R mediated **A to R** conversions<br>(only between neighboring nucleosomes) | $c_2 \sum_{i=1}^{128} R_i \frac{A_{i-1} + A_{i+1}}{2}$ |
| *Erasure* | Direct **R to A** conversions | $c_3 \sum_{i=1}^{128} R_i$ |
| *Stable Silencing* | Direct **R to I** conversions<br>(only at promoter) | $c_4 \sum_{i=63}^{67} R_i$ |

As explained above, all of the four reactions - nucleation, spreading, erasure and stably silencing - are defined as attempt rates, whereas the reaction propensities are defined as successful attempt rates. To be consistent with the success propensities, we simulated the model as a hybrid consisting of a first event-based gillespie update to select the attempted reaction (and to update the time-counter), as well as a subsequent reaction-type algorithm as an agent-based model by selecting nucleosomes as explained below. Simulations start with all 128 nucleosomes being in the A state. All reaction attempt propensities are defined and stored in a total attempt propensity vector *a* = [*c*_1_, 128*c*_2_, 128*c*_3_, 5*c*_4_] and the sum of all rates has been calculated and named as *a_total_* = *c*_1_ + 128*c*_2_ + 128*c*_3_ + 5*c*_4_. At each update step, the time is updated by generating a random number *ran*_1_ ∈ *uniform*(0,1) and then selecting a time interval *Δt* = 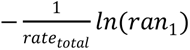. The selection of a reaction occurs at each update step, by first generating a random number *ran*_2_ ∈ *uniform*(0, *rate_total_*) and then choosing the reaction with attempt propensity *a_k_*, with k being the smallest value that fulfills 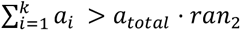. After the reaction attempt has been selected, one of the following reaction types is executed:

a) **Nucleation**: select the nucleosome at position 62 (the TetO position) and change its state to R if it is in A state.
b) **Spreading:** randomly select one nucleosome in the array from which the spreading will take place and, if it is in R state, choose one nearest neighboring nucleosome (right or left neighbor with equal probability) and change its state to R, if it is in A state.
c) **Erasure:** randomly select one nucleosome in the system and if it is in R state, change it to A.
d) **Stable silencing:** randomly select one nucleosome at positions 63 to 67 (the pEF promoter position) and change it to an I state, if it is in R state.

### Simulation of KRAB recruitment time courses

Simulation time courses were generated by running 1000 simulations as explained above for a total of 20 days and by recording the nucleosome state vector every 20 minutes in simulation time. The fraction of Citrine Off cells was determined at every recording by counting the number of inactive promoters (nucleosome state vectors with at least one R or I state at positions 63 to 67) and dividing it by the total number of simulations (1000). Nucleation rates with non-zero values were only present for a specified number of days (matching dox recruitment), before they were set to 0 ℎ^−1^ (dox release).

### Simulation of epigenetic memory landscapes

We performed simulations for 23 distinct durations of nucleation ranging from 0 to 11 days, in 0.5-day increments and we used 23 different nucleation rates for simulations, which varied logarithmically from 0.01 to 10. In total for both landscapes, we thus performed 529 (23×23) simulations for each unique pair of nucleation-rate and -duration, where we simulated 1,000 cells in parallel. The Z-axis in our representation displays the percentage of cells that were silenced by day 14 following nucleation, meaning that these cells were first simulated with the specified nucleation-rate and -duration before entering a 14-day simulation period devoid of any nucleation.

### Software

Flow cytometry data was analyzed using Matlab R2016a (Mathworks), FlowJo 10.9 (BD) and Easyflow ^92^. Fiji (ImageJ) was used to quantify western blot band intensities. Figures were generated using Graphpad Prism 9.5 (Dotmatics), Illustrator 2020/2023 (Adobe), and Biorender.

### Statistical analysis

All data are represented as mean ± SD or mean ± SEM unless otherwise stated. N indicates the number of independent experiments. Data was analyzed using a one-way ANOVA or Welch’s T-test when appropriate in GraphPad Prism 9.5. Statistical significance of the data is indicated as follows: *p ≤ 0.05, **p ≤ 0.01, ***p ≤ 0.001, ****p ≤ 0.0001; ns = not significant.

## Data availability

Illumina sequencing data generated in this study will be deposited to the NCBI GEO database.

## Code availability

Scripts used to analyze data and code for simulations will be available on Zenodo and Github.

## Material Availability

Plasmids generated in this study will be made available on Addgene. Additional materials may be available upon reasonable request from the lead contact.

## Acknowledgements

We thank James Ferrell, Joanna Wysocka, Or Gozani and Dylan Husmann (Stanford) for helpful discussions throughout the project. We also thank Howard Chang’s lab (Stanford) for use of their sonicator, Will Greenleaf’s lab (Stanford) for use of their MiSeq, and Mike Bassik’s lab (Stanford) for use of their NextSeq. We are grateful to Ben Doughty and Sage Allen for assistance with EMseq and Nicole DelRosso for sharing the rTetR-VP64 plasmid. We thank Josh Tycko, Ewa Bielczyk-Maczyńska and the members of the Bintu lab for comments on the manuscript. J.S. was supported by NIH Training Program grant T32-GM113854. A.R.T. was supported by the Sarafan Chem-H Chemistry-Biology Interface Training Grant and the NIH Training Program grant T32GM145402 This work was supported by the NIH-NIGMS MIRA R35GM12894701 and NIH 4D Nucleome grant U01DK127419 awarded to L.B.

## Author contributions

J.S and L.B conceived of project and experimental design. J.S engineered all cell lines related to the study and generated all flow cytometry time-course data unless otherwise stated. J.F.N generated the 3-state spreading model and performed simulations and experimental validation of simulations with flow cytometry. A.R.T and J.S performed ChIP and CUT and RUN experiments. C.H.L wrote the CUT & RUN analysis pipeline and performed formal analysis of genomics data. B.N.A performed experiments related to the activation of minCMV reporter and assessment of cell growth assays in HEK293T. J.S performed EMseq. A.R.T performed analysis of the EMseq data. M.M.H optimized the EMseq protocol and designed EMseq primers used in the study. J.S and J.W cloned plasmids used in the study. D.F generated the parental *H3F3B* edited TC28a2 cell lines used in the study. J.S and L.B wrote the manuscript with input from all authors.

---

**Fig. S1.**
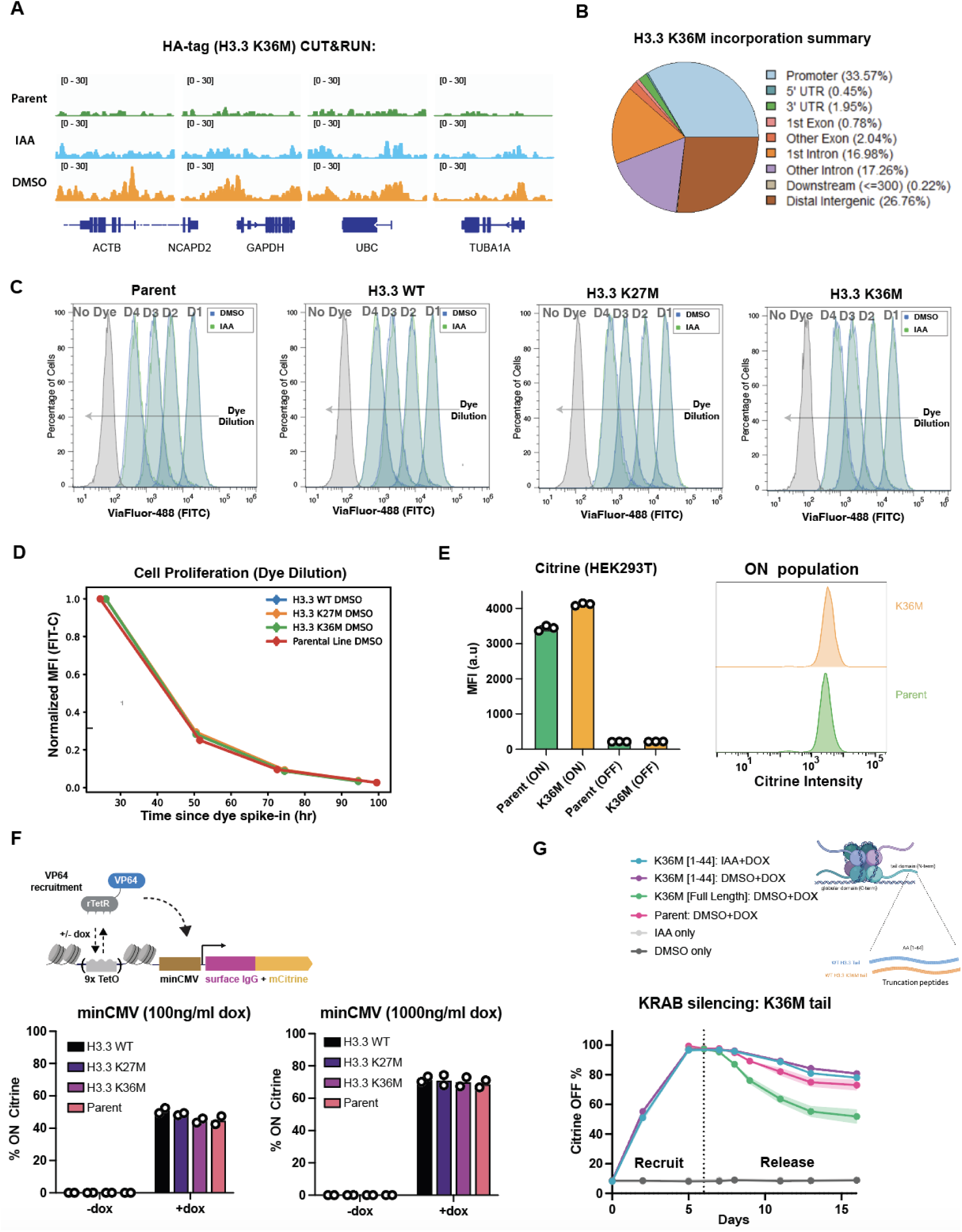
Characterization of the HEK293T reporter line. **(A)** CUT&RUN genome tracks against the HA-tag (H3.3K36M) in HEK293T cell cells after 48hr treatment with either 50μM IAA or DMSO. IAA dependent incorporation of H3.3 K36M is measured at different endogenous control genes. The parental (WT) cell line is included as a reference. Read numbers are normalized by E. coli spike-in. **(B)** Pie chart describing top enriched molecular function GO terms H3.3 K36M incorporation based on CUT&RUN data (HA-tag signal peaks). Dots are color codes by p-value and gene counts are represented by dot size and each color describes enrichment distribution of H3.3 K36M based on genetic element (e.g. promoter). **(C)** Flow cytometry time course tracking growth mediated dilution of Viafluor 488 SE cell proliferation dye in the FITC (Citrine) channel of WT HEK293T cells and HEK293T cells expressing WT, K27M, and K36M variants. Proliferation was measured in the presence of 50μM IAA or DMSO. **(D)** Quantification of mean fluorescence intensity (MFI) from flow cytometry distributions in **C** comparing the relative growth rates of HEK293T cell lines in the presence of DMSO (histone overexpression). Data from each cell line is normalized to the fluorescence intensity of the first time point. **(E)** Mean fluorescence intensity (MFI) of Citrine expression and representative flow cytometry distributions of basal Citrine reporter expression of parental HEK293T cells and HEK293T cells expressing H3.3 K36M. Background silenced cells (Citrine negative) are omitted from analysis of the ON population. **(F)** Quantification of transcriptional gene activation from a reporter driven by a minimal promoter (minCMV) upon dox mediated recruitment of rTetR-VP64 in the presence of oncohistone expression. Cells were treated with 100ng/ml dox (left) and 1000ng/ml dox (right). The % of activated Citrine positive cells were quantified by flow cytometry 48hr post dox addition. **(G)** Flow cytometry time course measuring silencing and memory after KRAB recruitment with 1000ng/ml in HEK293T cells expressing a truncated version of H3.3 K36M lacking the globular domain (only amino acids 1-44 of the N-terminal histone tail) to prevent nucleosomal incorporation (n=3 biological replicates). Shading represents mean +/- SEM. Dashed line represents the time at which the dox was washed out for memory time points.

**Fig. S2.**
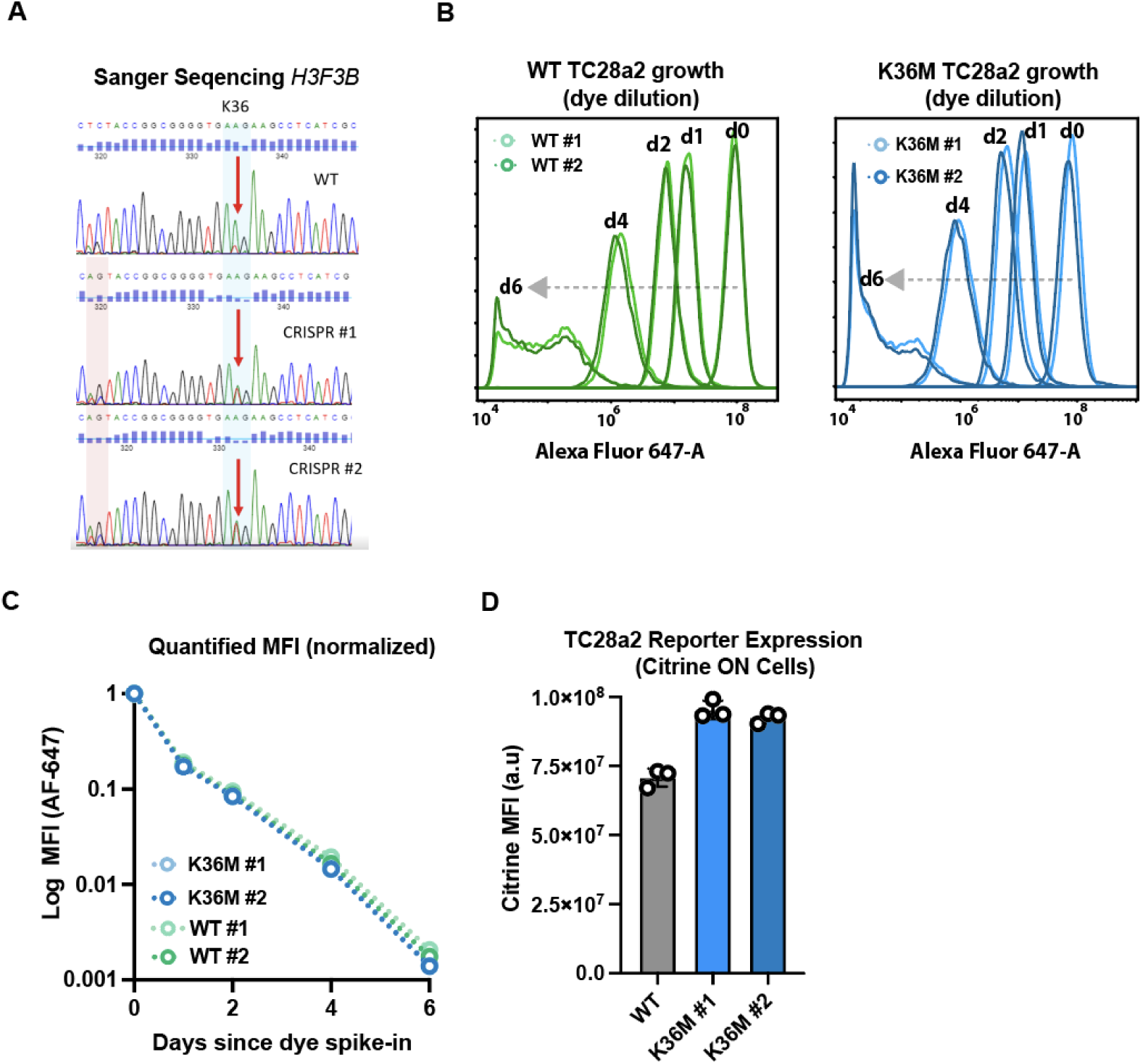
Characterization of TC28a2 reporter cell lines. **(A)** Sanger sequencing of CRISPR edited *H3F3B* TC28a2 clones showing the K36M point mutation. **(B)** Flow cytometry time courses tracking growth mediated dilution of CellTrace cell proliferation dye in the far-red channel. Compared are growth rates of WT TC28a2 cells (upper panel) and the two H3.3 K36M TC28a2 clones (lower panel). (**C)** The MFI of the dye used in **B** is quantified over time for each cell line to generate growth rate plots for WT and H3.3 K36M cells. **(D)** Bar plots measuring the MFI of the basal reporter expression in TC28a2 lines without any recruitment of KRAB. Background silenced cells (Citrine negative) are omitted from analysis.

**Fig. S3.**
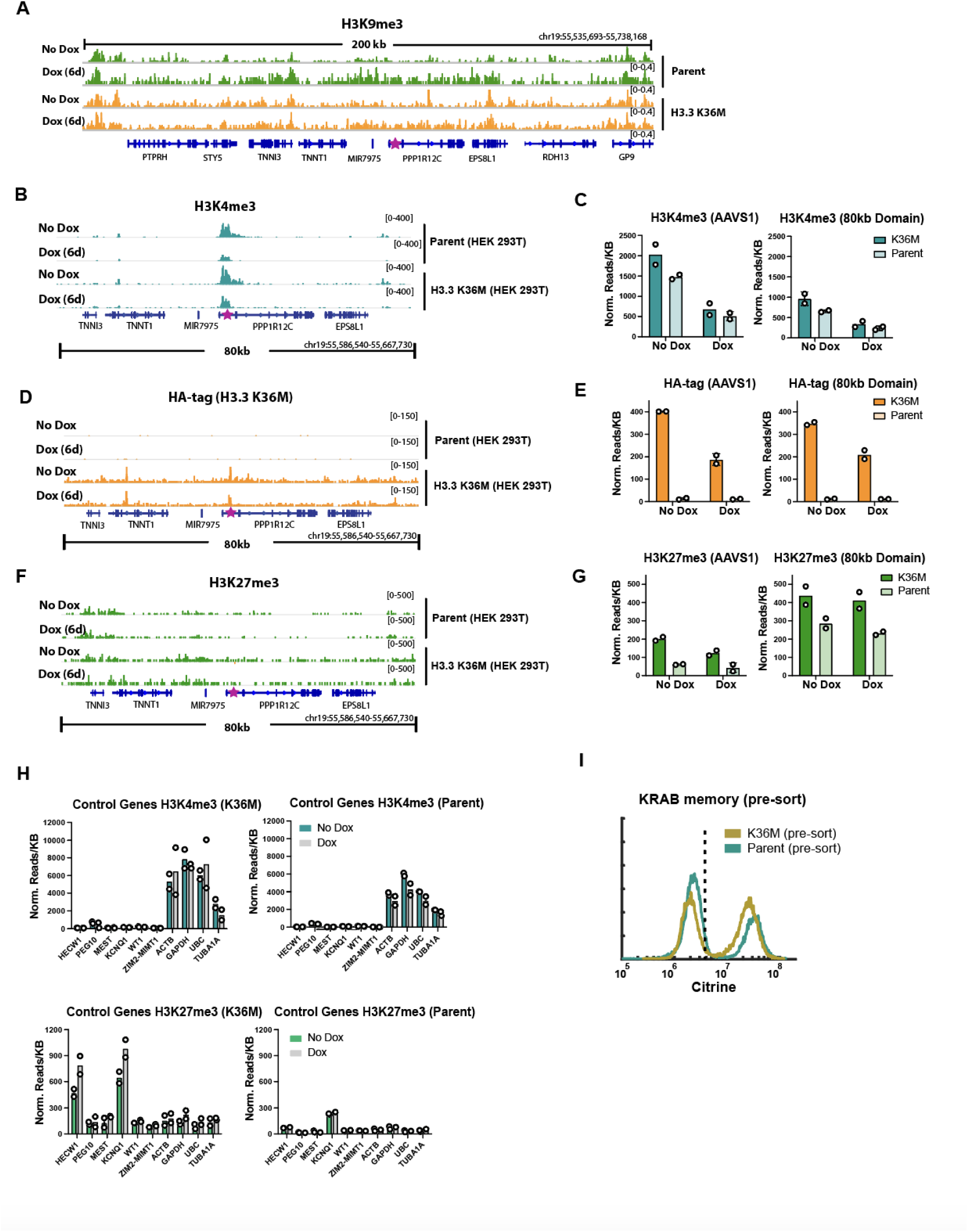
CUT&RUN of histone modifications at the reporter and control genes. **(A)** CUT&RUN genome tracks of an expanded 200kb region (from Fig. 3C) around the Citrine reporter measuring H3K9me3 in H3.3 K36M and parental lines after KRAB recruitment. The reporter integration site is demarcated with a star. Read numbers are CPM normalized. **(B)** CUT & RUN genome tracks of H3K4me3 in an 80kb domain around the reporter integration site within the *PPP1R12C* gene (AAVS1 locus) upon dox mediated KRAB recruitment (6 days) in both parental and H3.3 K36M HEK293T cells. Data is normalized using K-MetStat spike-in nucleosomes. The reporter integration site is demarcated with a star. **(C)** Quantification of normalized reads/kb of H3K4me3 signal shown in B at both the *PPP1R12C* gene (AAVS1 locus) and the entire 80kb domain (n=2 biological replicates). **(D)** CUT&RUN genome tracks of HA-tag (H3.3 K36M) incorporation in an 80kb domain around the reporter integration site within the *PPP1R12C* gene (AAVS1 locus) upon dox mediated KRAB recruitment (6 days) in H3.3 K36M HEK293T cells. The parental cell line is included as reference for background. Data is E. Coli spike-in normalized. The reporter integration site is demarcated with a star. **(E)** Quantification of normalized reads/kb of HA-tag signal shown in D at both the PPP1R12C gene (AAVS1 locus) and the entire 80kb domain (n=2 biological replicates). **(F)** CUT&RUN genome tracks of H3K27me3 an 80kb domain around the reporter integration site within the *PPP1R12C* gene (AAVS1 locus) upon dox mediated KRAB recruitment (6 days) in both parental and H3.3 K36M HEK293T cells. Data is normalized using K-MetStat spike-in nucleosomes. The reporter integration site is demarcated with a star. **(G)** Quantification of normalized reads/kb of H3K27me3 signal shown in F at both the *PPP1R12C* gene (AAVS1 locus) and the entire 80kb domain (n=2 biological replicates). **(H)** K-MET stat normalized H3K4me3 and H3K27me3 CUT&RUN signal in reads/kb quantified at a panel of endogenous genes within the presence and absence of dox mediated KRAB recruitment to the AAVS1 locus (6 days). **(I)** Flow cytometry distributions of Citrine reporter expression in H3.3 K36M and parental cells after establishment of KRAB memory (day 20) prior to magnetic cell separation for CUT & RUN.

**Fig. S4.**
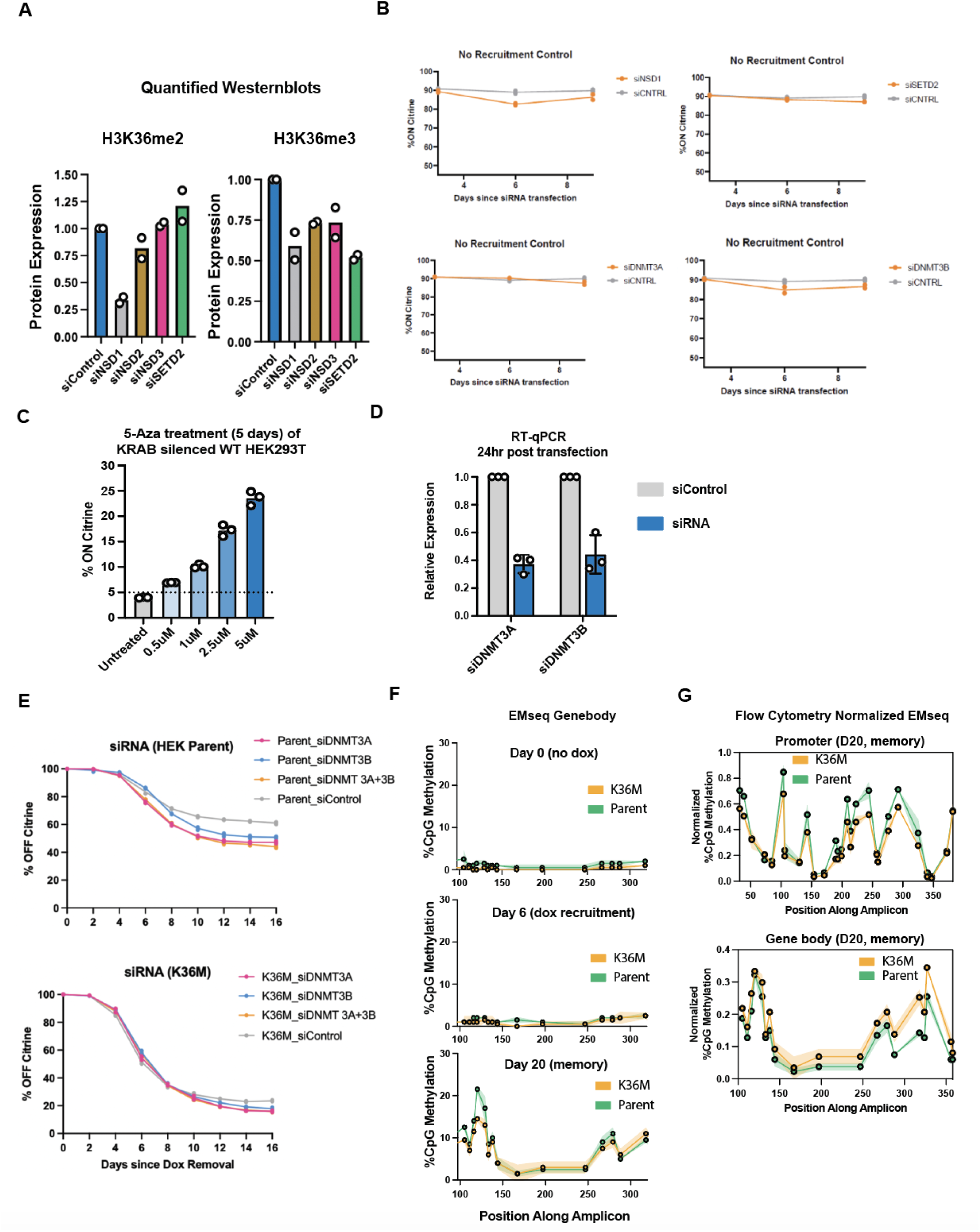
Validation of siRNA and DNA methylation effects in HEK293T cells. **(A)** Quantification of western blot band intensities measuring levels of H3K36me2 and H3K36me3 in HEK293T cells harvested after 48hr post transfection of siRNA against NSD family of methyltransferases and SETD2. (n=2 biological replicates). **(B)** Flow cytometry time course measuring levels of Citrine reporter expression in HEK293T cells transfected with siRNA targeting methyltransferases used in the study without recruitment of KRAB (n=2 biological replicates). **(C)** Quantification of % reporter reactivation in parental HEK293T after treatment with different concentrations of DNA methylation inhibitor 5-Aza-2’-deoxycytidine (5-Aza) for 5 days (n=3 biological replicates). Cells were first silenced with KRAB for 6 days. Stably silenced memory cells (14 days post dox release) were then enriched prior to addition of the 5-Aza for 5 days to induce reactivation. **(D)** RT-qPCR of DNMT3A and DNMT3B mRNA levels in in wildtype HEK293T cells 24hr post siRNA transfection. Data is normalized to non-targeting siControl transfection (n=3 biological replicates). **(E)** Flow cytometry time courses of epigenetic memory after 2 days of KRAB recruitment where siRNA targeting DNMT3A, DNMT3B, or a combination of DNMT3A/B was used upon dox removal (n=3 biological replicates). Memory was measured as %OFF Citrine every two days. **(F)** EMseq quantification of the %CpG methylation in the amplicon corresponding to the gene body of the reporter after KRAB recruitment and release (n=2 biological replicates). Data is matched with timepoints shown in Fig. 4 E&F. Shading represents standard deviation. **(F)** EMseq quantified %CpG methylation measured at the reporter region corresponding to the gene body after KRAB recruitment in parental and H3.3 K36M cells (n=2 biological replicates). **(G)** Normalized %CpG methylation at the promoter and gene body from the EMseq data shown in Fig. 4F and Fig. S4F (n=2 biological replicates). %CpG methylation is normalized by dividing the %CpG from EMseq by the corresponding % Citrine OFF cells in Fig. 4E which was used as the input for the assay. The memory (D20) timepoint was chosen for analysis due to sufficient levels of DNA methylation for comparison. Shading represents the standard deviation.

**Fig. S5.**
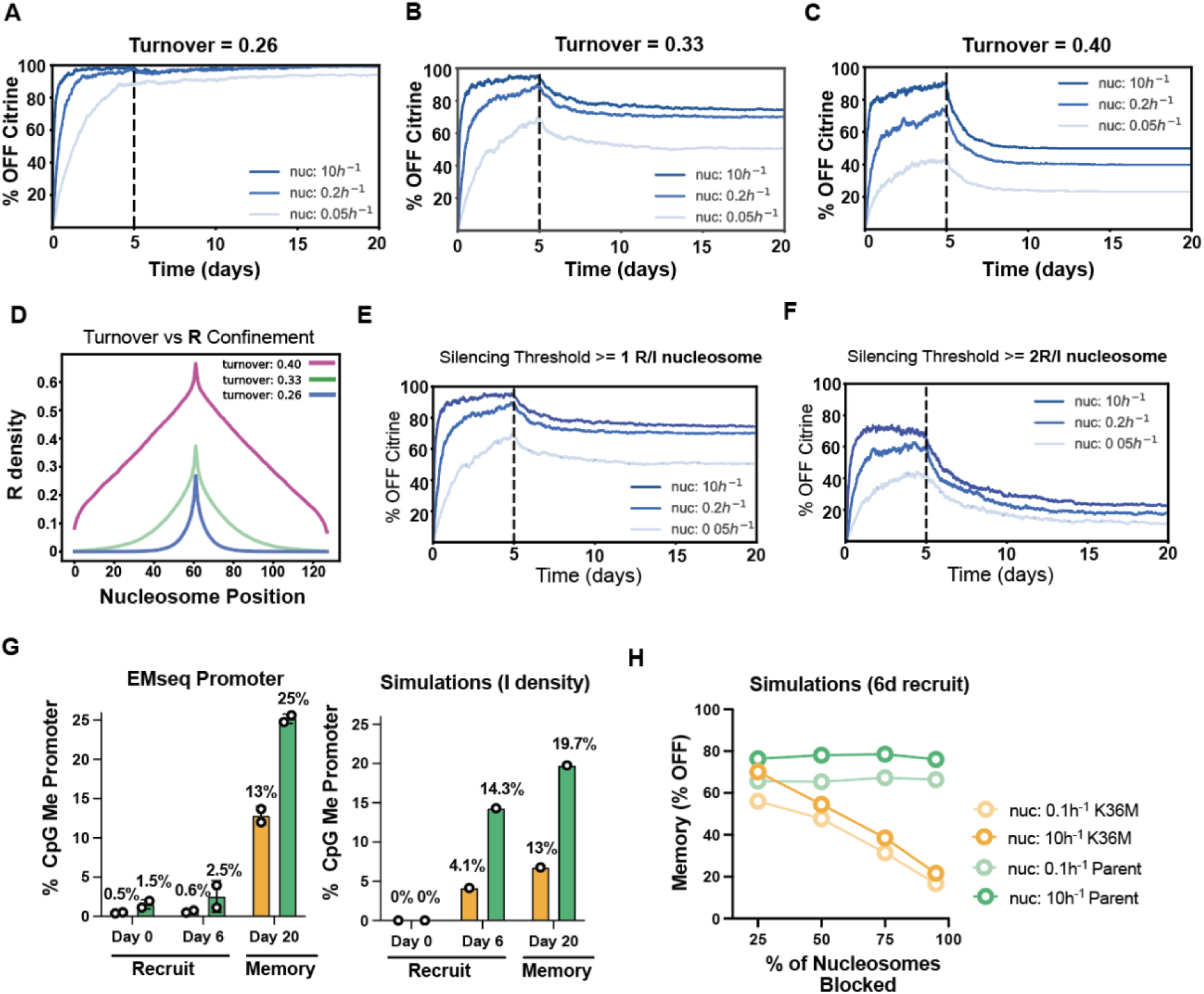
The model assumes a fine-tuned ratio of erasure and spreading rates, a silencing sensitive promoter, and predicts a reduced accumulation rate of DNA methylation in H3.3 K36M cells. We define promoter silencing when least ⅕ promoter nucleosomes are either in the R or I state (see methods). The model is based on an additional two major assumptions: (**A)** The spreading rate (chosen to be 1 per nucleosome per hour) must be precisely balanced with the erasure rate (chosen to be 0.33 per nucleosome per hour) such that the resulting ratio of 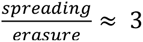. (**B)** If the ratio is lower, confinement is lost, and cells cannot reactivate, leading to ∼100% memory **(C)**. If the ratio is higher, cells are unable to be effectively silenced. **(D)** The system is very close to a critical point before loss of inherent confinement ^78,79^. (**E)** The promoter has to be very sensitive to R and I nucleosomes with one R or I nucleosome being sufficient to silence the whole promoter. **(F)** If we assume that more nucleosomes need to be in the repressed states to lead to gene silencing, we cannot get full promoter silencing. (**G)** The model predicts that the average percent of I nucleosomes accumulates more slowly in K36M cells compared to wt cells upon nucleation (left) in agreement with experiments (right). However, in contrast to DNA methylation density measured by EM-seq, I nucleosomes start to accumulate earlier (Day 6) and the difference in percent of I nucleosomes is larger in simulations than the measured difference in average percent of DNA methylation in experiments (n= 2 biological replicates). These differences could be explained by simplifications in the model such as the assumption that all promoter nucleosomes have the same rate of R to I transitions, whereas promoter DNA methylation is likely to depend on size and distribution of CpG islands ^93–95^. (**H)** Plot showing the percent of silent cells at day 14 of memory following 6 days of nucleation for parent cells (green) and K36M cells (orange). A nucleation rate of 0.1 ℎ^−1^ (lighter shade) and 10 ℎ^−1^(darker shade) are simulated. K36M memory landscapes and time course plots were generated using 75% of randomly selected nucleosomes being blocked for R to I transitions, because this percentage is in a similar range as suggested by measured fold changes in H3K36me2 and H3K36me3 abundance between K36M and Oncohistone cells measured by western blot and because this percentage was best in predicting the percent of permanently silenced cells in experiments.

